# A switch in cilia-mediated Hedgehog signaling controls muscle stem cell quiescence and cell cycle progression

**DOI:** 10.1101/2019.12.21.884601

**Authors:** Sara Betania Cruz-Migoni, Kamalliawati Mohd Imran, Aysha Wahid, Oisharja Rahman, James Briscoe, Anne-Gaëlle Borycki

## Abstract

Tissue homeostasis requires a tight control of stem cells to maintain quiescence in normal conditions, and ensure a balance between progenitor cell production and the need to preserve a stem cell pool in repair conditions. Using ex-vivo and in-vivo genetic approaches, we provide evidence that primary cilium-mediated repressive Hedgehog (Hh) signalling is required to maintain skeletal muscle stem cells (MuSCs) in a quiescent state. De-repression and further activation of Hh signalling initiates MuSC entry and progression through the cell cycle, and controls self-renewal to ensure efficient repair of injured muscles. We propose a model whereby disassembly of primary cilia upon MuSC activation induces a switch in Hh signalling from a repressive to active state that controls exit from quiescence. Positive Hh response in bi-potential muscle progenitor cells regulates also cell cycle progression and drives MuSC self-renewal. These findings identify Hh signalling as a major regulator of MuSC activity.

**Highlights:**

- Cilia-containing quiescent MuSCs are Hh signalling suppressed
- MuSC activation coincides with a switch to active Hh signalling
- *Smo* mutation delays cell cycle entry and progression, and causes impaired self-renewal
- *Ptch1* mutation promotes exit from quiescence, rapid cell cycle and increased self-renewal

**Graphical abstract:** 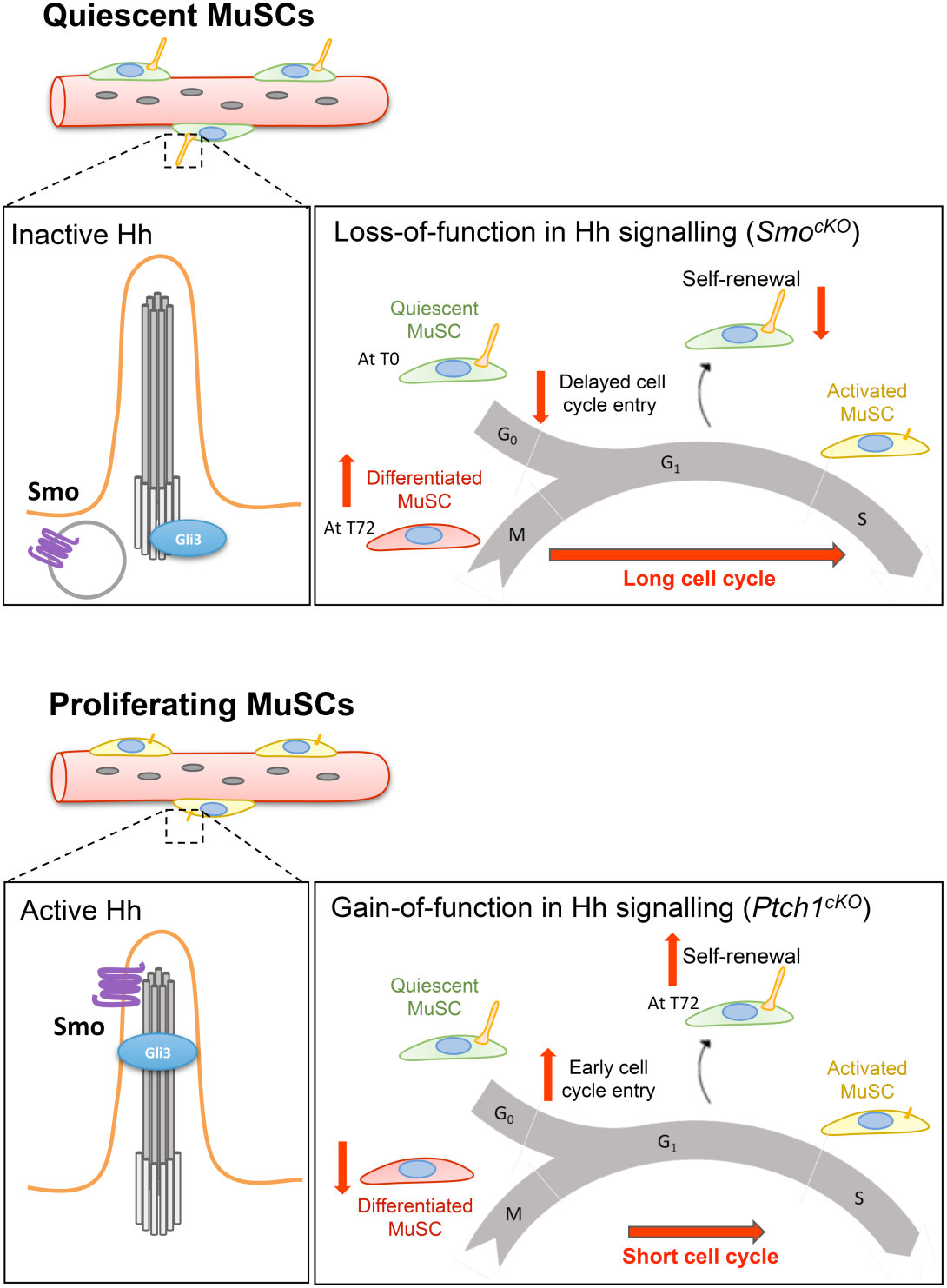

## Introduction

Skeletal muscle is one of a handful mammalian tissue capable of regeneration throughout the life course, largely due to the activity of skeletal muscle stem cells (MuSCs), characterized by the expression of the transcription factor Pax7 (Gunther et al., 2013; Lepper et al., 2011; Relaix and Zammit, 2012; Sambasivan et al., 2011; von Maltzahn et al., 2013). MuSCs originate from the embryonic dermomyotome, are deposited under the muscle fibre basal lamina during fetal development and are maintained in a reversible quiescent state in adult muscles (Gros et al., 2005; Kassar-Duchossoy et al., 2005; Relaix et al., 2005; Schienda et al., 2006). However, MuSCs can become activated in response to muscle injury, exercise or degeneration and exit G0 to enter the cell cycle and produce myogenic progenitor cells under the control of the myogenic regulatory factors Myf5 and MyoD (Dumont et al., 2015). Myogenic progenitor cell differentiation orchestrated by Myogenin then leads to the fusion of new myocytes to the damaged fibres, whereas down-regulation of MyoD and Myogenin and maintenance of Pax7 characterizes cells returning to quiescence in a self-renewal process essential for the long-term maintenance of a stem cell pool (Motohashi and Asakura, 2014). A tight regulation of MuSC quiescence and of the balance between proliferation, differentiation and self-renewal is crucial to ensure the maintenance of muscular integrity throughout life. Signalling molecules, in particular of the Wnt, Notch, HGF, IGF and TGFβ-related families, are key external cues previously reported to act cooperatively to preserve this equilibrium (Brack and Rando, 2012; Yin et al., 2013).

We recently reported that quiescent MuSCs harbour a primary cilium, a microtubule-containing organelle anchored at the cell surface via a modified centrosome called the basal body (Jaafar Marican et al., 2016). Notably, the primary cilium di-assembles upon MuSC activation and entry into the cell cycle, and re-assembles selectively at the surface of self-renewing, but not differentiating muscle progenitor cells upon cell cycle exit (Jaafar Marican et al., 2016). This raises the attractive possibility that Hedgehog (Hh) signalling plays important role in the control of MuSC activity, as the cellular machinery required for the transport, processing and activity of Hh pathway components, including the Gli transcription factors, is dependent on primary cilia (Briscoe and Therond, 2013).

In the absence of Hh ligand, Patched 1 (Ptch1), the receptor for Hh, prevents the transmembrane protein Smoothened (Smo) from localising to the primary cilium, causing GPR161-dependent activation of Protein Kinase A (PKA), which primes Gli3, and to a lesser extent Gli2, for further phosphorylation by Casein Kinase 1 (CK1) and Glycogen synthase kinase-3 beta (GSK3β), their association to Suppressor of Fused (SuFu) and proteolytic cleavage by the cilia-regulated proteasome located at the basal body into transcriptional repressors of Hh target genes (Corbit et al., 2005; Pal et al., 2016; Rohatgi et al., 2007; Wen et al., 2010). Hh binding to Ptch1 leads to its removal from cilia, the exit of Gpr161, and the accumulation and activation of Smo in cilia (Corbit et al., 2005; Pal et al., 2016). Activated Smo prevents also SuFu interaction with Gli2 and Gli3, resulting in the accumulation of Gli2 and Gli3 at the ciliary tip and their maturation into transcriptional activators (Chen et al., 2009; Humke et al., 2010). Consistent with a possible role for Hh signalling in adult skeletal muscles, previous studies have reported that blocking Hh signalling impeded muscle regeneration and conversely Hh treatment enhanced muscle regeneration in a mouse model of ischemia and muscular dystrophy (Palladino et al., 2012; Palladino et al., 2011; Piccioni et al., 2014a; Piccioni et al., 2014b). However, the cellular target(s) of Hh signalling has not been determined and it remains unclear whether Hh acts directly on MuSCs or not. Furthermore, the role of Hh signalling in adult muscles remains controversial with some reports claiming a pro-differentiating function and others reporting a trophic function (Elia et al., 2007; Fu et al., 2014; Koleva et al., 2005; Li et al., 2004), Given that primary cilia are required for maintaining Hh signalling repressed in the absence of ligand as well as for mediating Hh signalling activation, it is likely that blocking ciliogenesis may have different effects to inactivating or blocking Smo activation.

Here, using pharmacological and genetic approaches to cause loss-of-function and gain-of-function in Hh signalling we show that Hh signalling acts cell-autonomously on MuSCs to control their exit from quiescence, progression through the cell cycle, and self-renewal. Our findings support a model whereby the primary cilium acts as a gatekeeper controlling Hh-mediated transcriptional output and cell fate choice in MuSCs with important consequences for MuSC long-term regenerative capability muscle and homeostasis.

## Results

### Gli-mediated Hedgehog response is suppressed in quiescent MuSCs and induced upon MuSC activation in adult myogenesis

Although the treatment with exogenous Sonic hedgehog (Shh) of in vivo models of muscle injury, including ischemia, crush and cardiotoxin-mediated injury, has a beneficial effect on muscle regeneration (Piccioni et al., 2014a; Piccioni et al., 2014b), there is currently no clear evidence that Hedgehog signalling acts directly upon MuSCs. Instead, current reports suggest an indirect role of Hedgehog signals on the production of pro-angiogenic factors by interstitial fibroblasts or on immune cells (Piccioni et al., 2014a; Pola et al., 2003; Rajurkar et al., 2014; Renault et al., 2013). In vitro, both C2C12 and primary myoblasts up-regulate the Shh target genes *Ptch1* and *Gli1* following Shh-triggered stimulation or induced differentiation (Elia et al., 2007; Koleva et al., 2005; Li et al., 2004). This suggests that MuSCs may become responsive to Hedgehog signals during muscle regeneration.

To establish whether MuSCs are direct targets of Hh signalling and determine the kinetics of Hh response during adult myogenesis, we isolated single myofibres from 6 to 8-week old Tg(GBS-GFP) mice, which carry a GFP reporter gene under the control of a promoter containing concatemerized Gli binding sites (Balaskas et al., 2012), and cultured them ex vivo for 72 hours (Figure 1A). No GFP expression was detected in quiescent Pax7^+^ MuSCs on freshly isolated myofibres (0h, Figure 1B,C), confirming that quiescent MuSCs are refractory to Hh signalling. In contrast, approximately 55% of Pax7^+^ MuSCs were GFP-positive by 24h, when MuSCs become activated. By 72h, GFP expression was associated with Pax7^+^ and Pax7^-^ cells, indicating that Hh response occurs in differentiated (Pax7^-^), as well as proliferating and/or self-renewing (Pax7^+^) MuSCs (Figure 1B,C).

**Figure 1.**
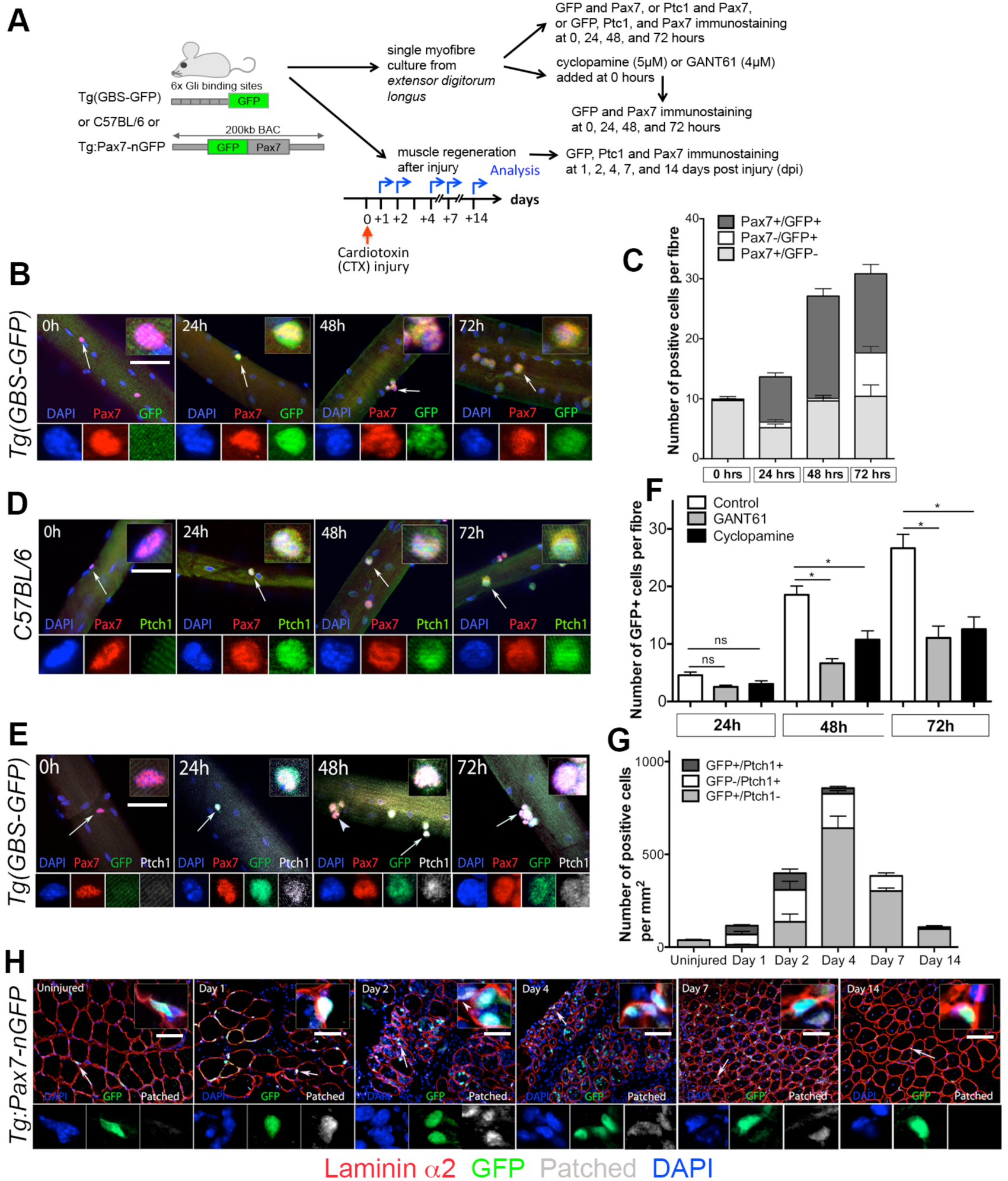
A switch in Hedgehog response coincides with MuSC exit from quiescence. (A) Schematic representation of experimental design. (B) Hedgehog response is observed in activated, but not quiescent MuSCs. Distribution of Pax7 (red) and GFP (green) in cultured Tg(GBS-GFP) myofibres. Magnified views for every channel are shown (regions indicated by an arrow). Scale bar: 50µm. n=3 with 25-55 fibres per time point. (C) Quantification of the Hedgehog response in cultured Tg(GBS-GFP) muscle fibres. Values are mean ± sem. (D) Distribution of Pax7 (red) and Ptch1 (green) in cultured C57BL/6 myofibres. Magnified views for every channel are shown (regions indicated by an arrow). n=3 with 35-55 fibres per time point. Scale bar: 50µm (E) Co-localization of GFP (green) and Ptch1 (white) in Pax7^+^ satellite cells (red) in cultured Tg(GBS-GFP) myofibres. Magnified views for every channel are shown (regions indicated by an arrow). Scale bar: 50µm (F) Quantification of Hedgehog response (number of GFP^+^ satellite cells) in Tg(GBS-GFP) myofibres cultured in the presence of GANT61 (4µM) or Cyclopamine (5µM). n= 3 with 25-30 fibres per time point. Values are mean ± sem. unpaired t test analysis *= p < 0.05. (G) Quantification of Ptch1 and GFP expression in Tg(Pax7-EGFP) muscles following injury. Values are mean ± sem. (H) Ptch1 distribution during muscle regeneration. Uninjured and injured Tg(Pax7-EGFP) muscles were analysed by immunofluorescence for GFP (green), Ptch1 (white) and Laminin alpha 2 (red). Inserts show high magnification images of cells indicated by white arrow. Individual channel images are shown below the main image. Scale bar: 50µm. See also Figure S1.

Consistent with these findings, Ptch1 was not detected in Pax7^+^ quiescent MuSCs at 0h, was up-regulated at 24h in Pax7^+^ activated MuSCs, and persisted as cells proliferated and differentiated in culture (48h and 72h, respectively; Figure 1D). Most cells expressing GFP were also positive for Ptch1, although some Ptch1^+^/GFP^-^ cells could be observed at 48h and 72h (Figure 1E). Thus, activated MuSCs are Hh responsive, and Ptch1 up-regulation is a more sensitive readout of Hh response than GFP in MuSCs. To confirm unequivocally that GFP detection marks Gli response in MuSCs, Tg(GBS-GFP) myofibres were cultured for 72h in the presence of cyclopamine, a Smo inhibitor (Cooper et al., 1998; Incardona et al., 1998) or GANT61, a small molecule inhibitor of Gli-mediated transcription (Lauth et al., 2007), and the distribution of GFP was examined. There was a significant reduction in the number of GFP-positive cells in myofibres cultured in the presence of Cyclopamine (40% at 48h and 53% at 72h) or GANT61 (60% at 48h and 58% at 72h) (Figure 1F and S1A), confirming that GFP expression occurs as the result of Hh signalling in MuSCs. Furthermore, as cyclopamine and GANT61 had similar effects on GFP expression, we concluded that Hh response in activated MuSCs occurs through the canonical Gli-dependent signalling pathway.

To assess Hh response in MuSCs in vivo, we carried out a unilateral cardiotoxin-mediated muscle injury of the *tibialis anterior* (TA) in Tg:Pax7-nGFP mice, a MuSC reporter line expressing EGFP under the control of the regulatory elements of Pax7 (Sambasivan et al., 2009) (Figure 1A). Ptch1 expression was assessed at 1, 2, 4, 7 and 14 days post injury (dpi) (Figure 1G,H). Ptch1 was undetectable in uninjured Tg:Pax7-nGFP muscles (Figure 1G,H), confirming that Hh signalling is inactive in quiescent MuSCs. At 1dpi, Ptch1 expression was observed in 80% of GFP^+^ cells (dpi) (Figure 1G,H). Although there was nearly a three-fold increase in the total number of GFP^+^ cells at 2dpi, the proportion of Ptch1^+^ MuSCs decreased to 40%, and decreased further to 4.5% at 4dpi (Figure 1G,H). By 7 and 14dpi, no GFP^+^ cell co-expressed Ptch1 (Figure 1G,H). Interestingly, we also observed Ptch1^+^ cells that were not labelled with GFP between 1-7dpi, indicating that Hh response occurs both in myogenic and non-myogenic cells following muscle injury, as reported previously (Pola et al., 2003; Rajurkar et al., 2014; Straface et al., 2009). From these experiments, we conclude that quiescent MuSCs are refractory to Hh signalling and become rapidly responsive to Hh signalling upon activation following muscle injury.

### Hedgehog signalling is required cell-autonomously for MuSC-mediated muscle regeneration

Given the switch in Hh response upon MuSC activation, we next asked whether Hh signalling played a role in MuSC activity and muscle regeneration. Mice carrying a conditional allele of *Smoothened* (*Smo^flox^*), an essential transmembrane transducer of Hh signalling, were bred with mice expressing an inducible Cre recombinase gene under the control of *Pax7* regulatory sequences (*Pax7^CreERT2^*) to generate *Pax7^CreERT2^,Smo^flox/flox^* mice (thereafter, named *Smo^cKO^*). Induction of recombination was initiated by four consecutive intra-peritoneal injections of tamoxifen, followed with a diet on tamoxifen chow. Consistent with previous reports (von Maltzahn et al., 2013), this regimen led to 83-90% recombination, as assessed by crossing *Smo^cKO^* mice into the *R26R-EYFP* background and examining EYFP^+^ cells in freshly isolated myofibres and in myofibres cultured for 24h (Figure S2A,B). Muscle injury was induced 3-5 days after the last tamoxifen injection by intramuscular injection of cardiotoxin, and muscles were analysed at 2, 4, 7, 14 and 21 days after injury (Figure 2A). Histological analyses showed that although some levels of regeneration occurred in *Smo^cKO^* mice, by and large *Smo^cKO^* muscles failed to repair efficiently as evidence by the presence of a large number of mononucleated cells and the higher frequency of fibres with a small cross-sectional area in *Smo^cKO^* compared to control (*Smo^flox/flox^*) injured muscles (Figure 2B-D and Figure S2C). Consistent with the absence of efficient regeneration, fibrosis measured by Collagen I distribution was prominent in *Smo^cKO^* compared to control injured muscles at 14dpi (Figure 2E,F). To characterize further the poor repair observed in *Smo^cKO^* mice, we determined the kinetics of muscle progenitor cell expansion by examining MyoD expression and the kinetics of differentiation by examining Myogenin expression between 2-14dpi (Figure 2G,H and Figure S2D,E). There was a delay in MyoD and Myogenin up-regulation in *Smo^cKO^* compared to control muscles at 2dpi (Figure 2G,H), suggesting that MuSCs ability to enter and/or progress through the myogenic program was impaired. Although there was no significant difference in the number of MyoD^+^ cells between *Smo^cKO^* and control muscles by 4 and 7dpi, we observed a 3.5-fold increase in the number of MyoD^+^ progenitor cells in *Smo^cKO^* compared to control muscles at 14 dpi (Figure 2G). A similar trend was observed with Myogenin resulting in a 16-fold increase in Myogenin^+^ cells at 14dpi in *Smo^cKO^* compared to control muscles (Figure 2H). This suggests that in the absence of Hh signalling muscle regeneration was delayed leading to the persistence of activated and differentiating muscle progenitor cells at 14dpi. To test this possibility, we next assayed the regeneration process at 21dpi. This revealed that muscle regeneration was not further advanced by 21dpi, as indicated by the high number of MyoD^+^ progenitor cells in *Smo^cKO^* mice compared to the restricted number of MyoD^+^ progenitor cells in control mice (Figure 2I). Therefore, we concluded that *Smo^cKO^* muscles fail to repair efficiently, and that signalling through Smoothened is essential for MuSC-mediated regeneration of skeletal muscles.

**Figure 2.**
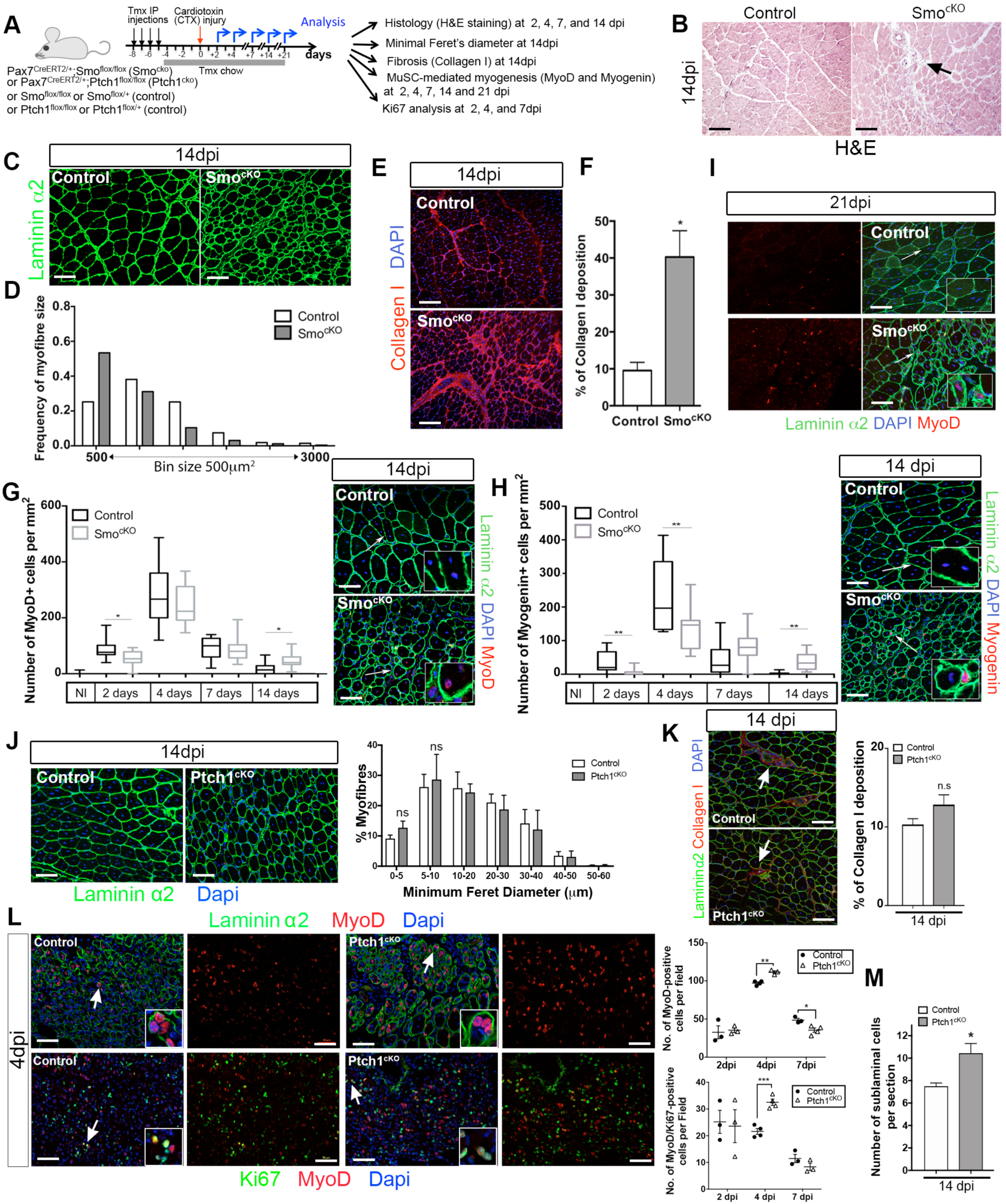
Levels of Hedgehog signalling control MuSC activity and muscle regeneration. (A) Schematic representation of experimental design. (B) Representative images of 14dpi *Tibialis anterior* muscle transverse sections from control and *Smo^cKO^* mice analysed by histology (H&E staining). Black arrow indicates infiltrating cells. Scale bar: 100µm. (C) Representative images of 14dpi *Tibialis anterior* muscle transverse sections from control and *Smo^cKO^* mice analysed by immunofluorescence using antibodies against Laminin α2 (green). Scale bar: 50µm. (D) Graph showing the distribution of myofibre cross-sectional area in 14dpi control (white bars) and *Smo^cKO^* (grey bars) *Tibialis anterior* muscles. n= 3 mice per genotype. (E) Representative images of 14dpi *Tibialis anterior* muscle transverse sections from control and *Smo^cKO^* mice analysed by immunofluorescence using antibodies against Collagen I (red). Nuclei are stained with Dapi (blue). Scale bar: 50µm. (F) Quantification of % Collagen I deposition in 14dpi *Tibialis anterior* muscle transverse sections from control (white bars) and *Smo^cKO^* (grey bars) mice. Values are mean ± sem. unpaired t test analysis *= p < 0.05. n= 3 mice per genotype. (G) Left: Graph showing the number of MyoD-positive cells per mm^2^ in control (black boxes) and *Smo^cKO^* (grey boxes) *Tibialis anterior* muscle transverse sections during regeneration following injury. Values are mean ± sem. unpaired t test analysis *= p < 0.05. n= 3 mice per genotype. Right: representative images of 14dpi *Tibialis anterior* muscle transverse sections from control and *Smo^cKO^* mice analysed by immunofluorescence using antibodies against MyoD (red) and Laminin α2 (green). Nuclei are stained with Dapi (blue). White arrows indicate the areas magnified in inserts. Scale bar: 50µm. (H) Left: Graph showing the number of Myogenin-positive cells per mm^2^ in control (black boxes) and *Smo^cKO^* (grey boxes) *Tibialis anterior* muscle transverse sections during regeneration following injury. Values are mean ± sem. unpaired t test analysis **= p < 0.01. n= 3 mice per genotype. Right: representative images of 14dpi *Tibialis anterior* muscle transverse sections from control and *Smo^cKO^* mice analysed by immunofluorescence using antibodies against Myogenin (red) and Laminin α2 (green). Nuclei are stained with Dapi (blue). White arrows indicate the areas magnified in inserts. Scale bar: 50µm. (I) Representative images of 21dpi *Tibialis anterior* muscle transverse sections from control and *Smo^cKO^* mice analysed by immunofluorescence using antibodies against MyoD (red) and Laminin α2 (green). Nuclei are stained with Dapi (blue). Left panels show the red channel (MyoD), and right panels show the merge images. White arrows indicate the areas magnified in inserts. Scale bar: 50µm. (J) Left: Representative images of 14dpi *Tibialis anterior* muscle transverse sections from control and *Ptch1^cKO^* mice analysed by immunofluorescence using antibodies against Laminin α2 (green). Nuclei are counterstained with Dapi (blue). Scale bar: 50µm. Right: Graph showing the distribution of myofibre minimum feret’s diameter (µm) in 14dpi control (white bars) and *Ptch1^cKO^* (grey bars) *Tibialis anterior* muscles. n= 3-4 mice per genotype. Values are mean ± sem. unpaired t test analysis *= p< 0.05, **= p < 0.01. (K) Left: Representative images of 14dpi *Tibialis anterior* muscle transverse sections from control and *Ptch1^cKO^* mice analysed by immunofluorescence using anitbodies against Collagen I (red) and Laminin α2 (green). Nuclei are counterstained with Dapi (blue). Scale bar: 50µm. Right: Percentage of Collagen I deposition per transverse section in 14dpi control and *Ptch1^cKO^* muscles. Values are mean ± sem. unpaired t test analysis. ns= non significant. (L) Left: Representative images of 4dpi *Tibialis anterior* muscle transverse sections from control and *Ptch1^cKO^* mice analysed by immunofluorescence using antibodies against Laminin α2 (green) and MyoD (red) (top panels), and MyoD (red) and Ki67 (green) (bottom panels). Nuclei are counterstained with Dapi (blue). Scale bar: 50µm. Right: Graphs showing the quantification of the number of MyoD-positive cells at 2, 4, and 7dpi (top graph), and the number of Ki67/MyoD-positive cells at 2, 4, and 7dpi (bottom graph) in control (black circles) and *Ptch1^cKO^* (open triangles) *Tibialis anterior* muscles. n= 3-4 mice per genotype per time point. Values are mean ± sem. unpaired t test analysis **= p < 0.01, ***= p< 0.001. (M) Graph showing the number of MuSCs returning in a sub-laminal position in 14dpi *tibialis anterior* muscles in control (white) and *Ptch1^cKO^* (grey) mice. Values are mean ± sem. unpaired t test analysis **= p < 0.01. See also Figure S2.

### Hh signalling gain-of-function mutation in MuSCs causes an increase in the number of muscle progenitor cells and self-renewing MuSCs

We next assessed the effect of up-regulating Hh signalling on skeletal muscle regeneration by crossing *Pax7^CreERT2^* mice to *Ptch1^flox/flox^* mice to generate *Pax7^CreERT2/+^*;*Ptch1^flox/flox^* mice (thereafter, named *Ptch1^cKO^)* (Figure 2A). After a single cardiotoxin-mediated muscle injury of the *tibialis anterior* muscle, we observed that muscle regeneration occurred nearly normally in *Ptch1*-deficient muscles (Figure 2J). In line with this efficient regeneration, there was no increased fibrosis in *Ptch1^cKO^* muscles (Figure 2K). Interestingly, regenerating *Ptch1^cKO^* muscles presented a transient increase at 4dpi in the number of MyoD-positive muscle progenitor cells (Figure 2L and S2F). As the increase of muscle progenitor cells correlated with an increased number of double labelled Ki67/MyoD cells in regenerating *Ptch1^cKO^* muscles (Figure 2L and S2G), we concluded that up-regulating Hh signalling promoted the proliferation of MuSC in a cell-autonomous manner. Furthermore, the reduced number of both MyoD-positive and Ki67/MyoD-positive cells at 7dpi in *Ptch1^cKO^* compared to control muscles suggested that *Ptch1^cKO^* MuSCs experience a shorter period of active proliferation, resulting in a smaller pool of muscle progenitor cells by 7dpi. Consequently, although the number of Myogenin-positive cells increased transiently at 4dpi, there was an overall reduction of the number of differentiating muscle progenitor cells at 7dpi in *Ptch1^cKO^* compared to control muscles (Figure S2H,I), consistent with the presence of a higher number of smaller fibres in *Ptch1^cKO^* compared to control muscles (Fiure 2J). Instead, regenerating *Ptch1^cKO^* muscles displayed a higher number of self-renewing stem cells returning in a sub-laminal position at 14dpi compared to control muscles (Figure 2M). Thus, up-regulation of Hh signalling caused a transient increase in muscle progenitor cell proliferation, and a concomitant increase in self-renewing MuSCs returning to quiescence and decrease in differentiating muscle progenitor cells.

### Levels of Hh signalling determine how MuSCs progress through adult myogenesis and self-renew

To determine the role of Hedgehog signalling in MuSC, we treated myofibre cultures with GANT61 to block Gli-mediated Hh signalling or with SAG (Smo agonist) to up-regulate Hh signalling (Figure 3A). GANT61 treatment caused an overall reduction in the number of MuSCs, whereas SAG treatment resulted in an increase in the number of MuSCs compared to control cultures treated with DMSO at 48 and 72h (Figure 3B,C). Immunofluorescence analyses showed that the cell populations most affected by the inhibition or up-regulation of Hh signalling were proliferating muscle progenitor cells (Pax7^+^/MyoD^+^) at 48h and 72h (Figure 3B,C). In addition, there was a significant decrease in the number of Caveolin^+^/Myogenin^-^ cells, which comprise proliferating progenitor cells and self-renewing cells, in the presence of GANT61 (Figure 3B,C). Conversely, the number of Caveolin^+^/Myogenin^-^ cells increased in the presence of SAG (Figure 3B,C). Treating myofibres with GANT61 for 60 hours, a time at which MuSC expansion slows down and differentiating progenitor cells emerge, revealed that in the presence of GANT61, a reduced proportion of cells had entered differentiation (Pax7^-^/MyoD^+^) and a higher proportion of cells remained proliferative (Pax7^+^/MyoD^+^) compared to control conditions (Figure 3D,E). The reduction in the number of Pax7^+^/MyoD^-^ cells at 60h and 72h (Figure 3E) suggested that loss of Hh signalling affected also self-renewal, an observation corroborated by the converse increase in the proportion of Pax7^+^/MyoD^-^ self-renewing cells upon Hh up-regulation (Figure S3A). By 72 hours, the ratio of differentiating (Pax7^-^ MyoD^+^) versus proliferating (Pax7^+^MyoD^+^) progenitor cells was noticeably affected with a far greater proportion of cells that had not progressed through the myogenic program (Figure 3D,E). Further analyses confirmed the general delay in the adult myogenic program, as fewer Pax7^-^Myf5^+^ cells, indicating the progression of progenitor cells from activated to differentiating cells, were observed in the presence of GANT61 compared to control cultures at 60 hours (Figure 3F and S3B). Likewise, immunofluorescence analysis of 72h-cultured myofibres with anti-MyoD and anti-Myogenin antibodies showed an absence of terminally differentiated cells (MyoD^-^/Myogenin^+^), a reduction in the number of differentiating myoblasts (MyoD^+^/Myogenin^+^), and a 2-fold increase in the number of proliferating myoblasts (MyoD^+^/Myogenin^-^) in the presence of GANT61 (Figure 3G and S3C). In contrast, stimulation of Hh signalling with SAG resulted in an increased number of differentiating muscle progenitor cells (Figure 3G and S3C). Taken together, these observations indicate that MuSC progression through the myogenic program is delayed in the absence of Hh signalling. Conversely, there is a greater proportion of muscle progenitor cells generated when Hh signalling is up-regulated. Our data indicate also that MuSC self-renewal is influenced by levels of Hh signalling.

**Figure 3.**
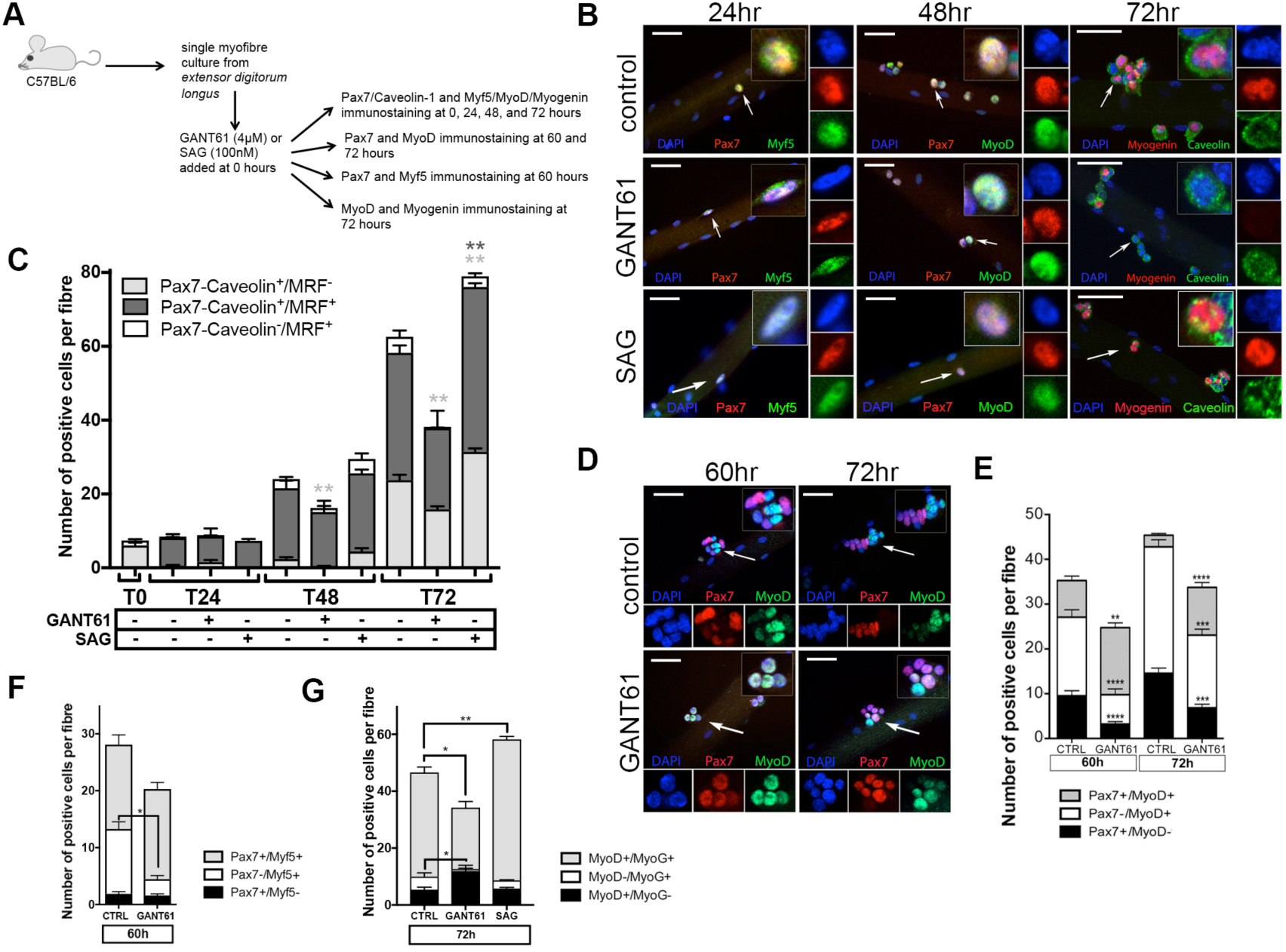
Hedgehog signalling is required for MuSC progression through the myogenic program. (A) Schematic representation of experimental design. (B) Immunofluorescence analysis of *extensor digitorum longus* myofibres cultured in the presence of DMSO (control), GANT61 (4µM), or SAG (100nM) using antibodies against Pax7 (red) and Myf5 (green) at 24h, Pax7 (red) and MyoD (green) at 48h and Myogenin (red) and Caveolin-1 (green) at 72h. White arrows indicate cells shown in insert (magnified view of merge image). Blue (Dapi), red (Pax7 or Myogenin) and green (Myf5 or MyoD or Cavelolin-1) channels are shown on right panels. Scale bar: 50µm. (C) Quantification of immunostaining shown in B. Values are mean ± sem, n= 3 experiments with 15-20 myofibres analysed per culture condition per time point in each experiment. unpaired t test analysis *= p < 0.05, **= p < 0.01. (D) Immunofluorescence analysis of C57BL/6 myofibres cultured in the presence of DMSO (control) or GANT61 (4µM) for 60 and 72h, using antibodies against Pax7 (red) and MyoD (green). White arrows indicate cells shown in insert (magnified view of merge image). Blue (Dapi), red (Pax7) and green (MyoD) channels are shown below the main image. Scale bar: 50µm. (E) Quantification of immunostaining shown in D. Values are mean ± sem. n=3 experiments with a total of 37-55 myofibres per time point per culture condition analysed. unpaired t test analysis. **= p < 0.01, ***= p < 0.005; ****= p < 0.001. (F) Quantification of immunofluorescence analysis using antibodies against Pax7 and Myf5 of C57BL/6 myofibres cultured in the presence of DMSO (control) or GANT61 (4µM) for 60h. Values are mean ± sem. n= 3 experiments with a total of 40-46 myofibres per culture condition analysed. unpaired t test analysis *= p < 0.05. (G) Quantification of immunofluorescence analysis using antibodies against MyoD and Myogenin of C57BL/6 myofibres cultured in the presence of DMSO (control), GANT61 (4µM), or SAG (100nM) for 72h. Values are mean ± sem. n=3 experiments with a total of 44-46 myofibres per culture condition. unpaired t test analysis *= p < 0.05, **= p < 0.01. See also Figure S3.

### Levels of Hh signalling control MuSC entry and progression through the cell cycle

Given our observations that levels of Hh signalling determine how MuSC progress through the myogenic program and the number of muscle progenitor cells generated, we tested whether levels of Hh signalling had an effect on MuSC proliferation (Figure 4A). The number of MuSCs positive for Ki67 in cultured myofibres treated with GANT61, SAG or DMSO was determined. After 48 and 72hr culture in the presence of GANT61, we observed a significant decrease in the number of Pax7^+^/Ki67^+^ and Pax7^-^/Ki67^+^ cells, corresponding to proliferating (MyoD^+^) and differentiating (Myogenin^+^) progenitor cells, respectively, compared to control conditions (Figure 4B,C). Although SAG treatment did not have a significant effect on MuSC at 48h, it caused an increase in the number of Pax7^+^/Ki67^+^ muscle progenitor cells at 72h (Figure 4B,C). Therefore, levels of Hh signalling are critical for the control of proliferation and the production of muscle progenitor cells. Additionally, as Ki67 marks all phases but G0 of the cell cycle (Scholzen and Gerdes, 2000), the reduced number of Pax7^+^/Ki67^-^ cells at 72h in GANT61-treated cultures (Figure 4B), confirmed that blocking Hh signalling impacts on MuSC ability to self-renew.

**Figure 4.**
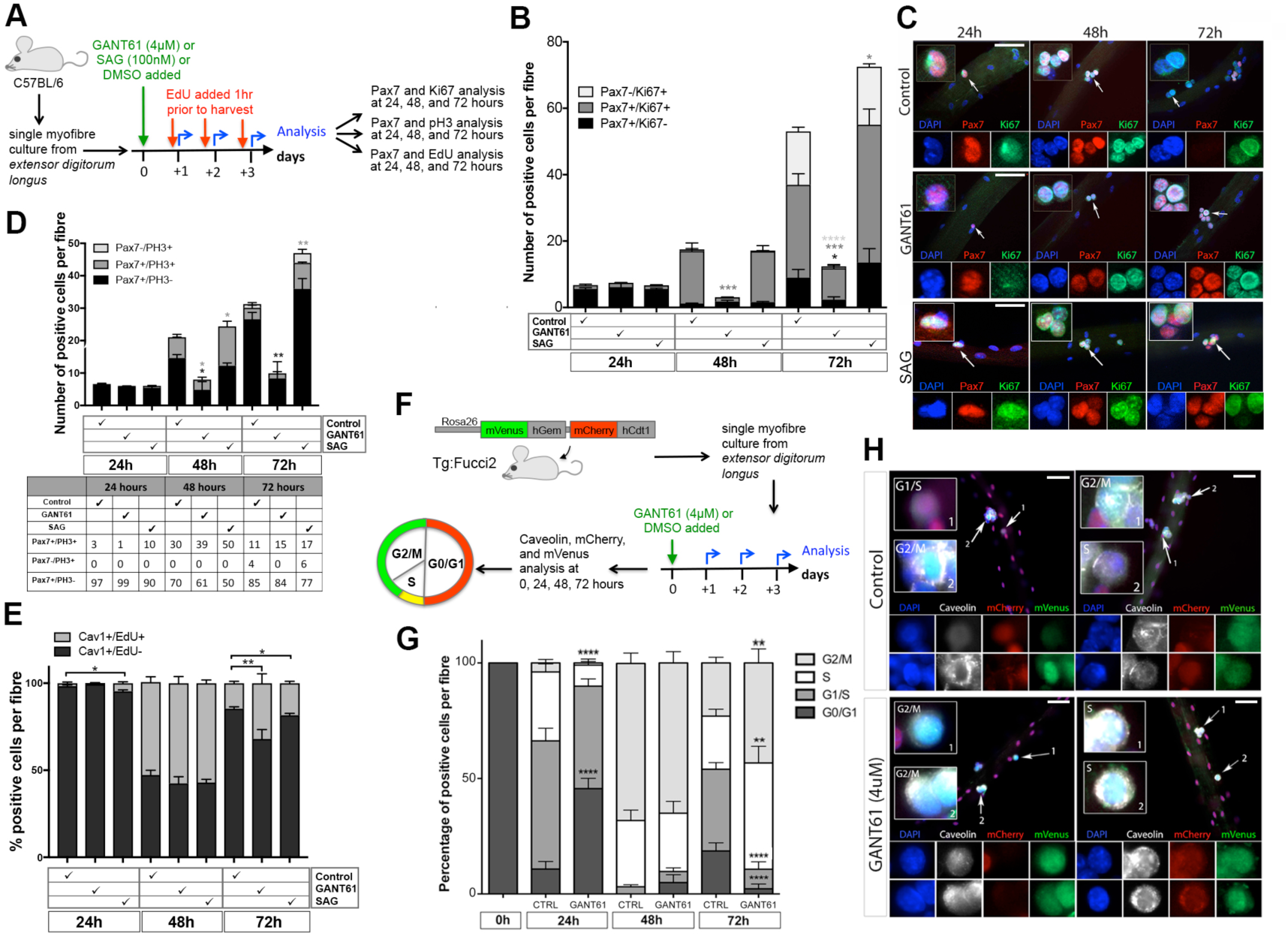
Hedgehog signalling controls MuSC progression through the cell cycle. (A) Schematic representation of the experimental design used for (B-E). (B) Quantification of the effect of increasing or decreasing Hh signalling on MuSC proliferation. Graph showing the number of positive cells per myofibre following immunofluorescence analysis using antibodies against Pax7 and Ki67 of C57BL/6 myofibres cultured in the presence of DMSO, GANT61 (4uM), or SAG (100nM). Values are mean ± sem. n=3 experiments with a total of 38-64 myofibres per time point per culture condition. unpaired t test analysis *= p < 0.05, **= p < 0.01, ***= p < 0.005; ****= p < 0.001. (C) Representative images of C57BL/6 myofibres analysed by immunofluorescence using antibodies against Pax7 (red) and Ki67 (green) at 24h, 48h, and 72h after culture in the presence of DMSO (control), GANT61 (4uM), or SAG (100nM). White arrows indicate cells shown in insert (magnified view of merge image). Blue (Dapi), red (Pax7) and green (Ki67) channels are shown below the main image. Scale bar: 50µm. (D) Top panel: Graph showing the number of MuSCs per myofibre in M phase. C57BL/6 myofibres were cultured in the presence of DMSO (control), GANT61 (4uM), or SAG (100nM) and analysed by immunostaining using antibodies against Pax7 and pH3. Values are mean ± sem. n=3 experiments with a total of 31-53 myofibres per time point per culture condition. unpaired t test analysis *= p < 0.05, **= p < 0.01. Bottom panel: Table showing the % of positive cell population. (E) Quantification of the number of MuSCs in S phase. C57BL/6 myofibres were cultured in the presence of DMSO (control), GANT61 (4uM), or SAG (100nM) for the indicated time, and analysed by immunostaining of EdU and Pax7 Values are mean ± sem. n=3 experiments with >49 myofibres per time point and culture condition. unpaired t test analysis *= p < 0.05, **= p < 0.01. (F) Schematic representation of the experimental design used in G and H, showing the expected colour combination of reporter gene expression during the cell cycle. (G) Graph representing the percentage of MuSC in each phase of the cell cycle as determined by immunofluorescence in control and GANT61-treated Tg:Fucci2 myofibres. Data is from three independent experiments with 25 to 35 myofibres for each time point. Values are mean ± sem. unpaired t test analysis *= p < 0.05, **= p < 0.01, ***= p < 0.005; ****= p < 0.001. (H) Representative immunofluorescence images of Tg:Fucci2 control and GANT61-treated myofibres analysed using antibodies against Caveolin-1 (white) and GFP (green). mCherry was measured directly. White arrows indicate cells shown with high magnification images in insert. Separate colour channels are shown below the main image. Scale bar: 50µm. See also Figure S4.

To further understand how Hh signalling controls MuSC proliferation, we examined the distribution of phospho-Histone 3 (pH3), which labels cells undergoing mitosis (Hans and Dimitrov, 2001). Consistent with our observations that levels of Hh signalling determine how MuSC progress through the myogenic program (Figure 3), we observed that blocking Hh signalling caused initially a reduction in the proportion of cells in M phase (1% in GANT61-treated versus 3% in control cultures at 24hr) (Figure 4D and S4A). However, although the total number of MuSCs per myofibre was reduced in the presence of GANT61, the percentage of cells in M phase was greater in GANT61-treated myofibre cultures compared to control cultures at 48h and 72h (Figure 4D and S4A). In contrast, up-regulation of Hh signalling consistently resulted in an increased number (and proportion) of cells in M phase throughout the culture period (10%, 50%, and 23% in SAG-treated versus 3%, 30% and 15% in control cultures at 24h, 48h, and 72h respectively) (Figure 4D and S4A). These findings suggested that levels of Hh signalling control MuSC rate of entry and progression through the cell cycle.

We next performed EdU pulse labelling in myofibre cultures to measure the proportion of cells entering S phase in the presence of GANT61 or SAG compared to control cultures. Confirming our previous findings, blocking Hh signalling caused a delay in MuSCs entering S phase whereas up-regulating Hh signalling had the converse effect and resulted in a higher proportion of MuSCs in S phase at 24h (Figure 4E). By 72hr, the proportion of muscle progenitor cells in S phase was greater in GANT61-treated compared to DMSO-treated myofibres (Figure 4E and S4B). Likewise, there was a significant increase in the percentage of muscle progenitor cells in S phase in SAG-treated myofibres compared to control myofibres (Figure 4E and S4B). Taken together, these observations indicate that loss of Hh signalling delays MuSC entry into the cell cycle, whereas up-regulation of Hh signalling drives MuSCs into the cell cycle. In addition, our data reveal that loss of Hh signalling affects further MuSC cell cycle progression once activated. In contrast, up-regulation of Hh signalling causes a steady increase in progenitor cell numbers as a result of a higher proportion of activated MuSCs progressing through the cell cycle.

To gain further understanding of the impact of blocking Hh signalling on MuSC cell cycle progression, we isolated and cultured myofibres from R26p-Fucci2 mice (thereafter named *Tg:Fucci2*), which carry a transgene containing mCherry-hCdt1 and mVenus-hGem under the control of the ubiquitous Rosa26 promoter (Abe et al., 2013). The ubiquitinylation domain of the human Cdt1 protein drives accumulation of mCherry (red) in the cell nucleus during G0, G1, and early S phases, while the ubiquitinylation of the human Geminin protein drives accumulation of mVenus (green) in S, G2, and M phases (Figure 4F and S4C). R26p-Fucci2 myofibres were cultured for 72h in the presence of GANT61, and green and red fluorescence was recorded to assess the proportion of cells in each phase of the cell cycle (Figure 4F-H and S4C). In freshly isolated myofibres, all myonuclei and 97.4% of Pax7^+^ cells were red confirming their quiescent state (G0). In agreement with our EdU incorporation data, fewer MuSCs entered and progressed through the cell cycle after 24hr in the presence of GANT61 compared to control (44% cells remained in G0/G1 compared to 9% in control condition). By 48hr, most MuSCs had entered the cell cycle, although some cells persisted in G0/G1 in the presence of GANT61. By 72hr when control muscle progenitor cells began exiting the cell cycle to differentiate, GANT61-treated muscle progenitor cells were accumulating in S phase, and to a lesser extent in G2/M (50% cells in S phase compared to 23% in control condition) (Figure 4G,H). Thus, in the absence of Hh signalling, MuSCs delay their entry into the cell cycle, and muscle progenitor cells appear to cycle at a slower pace and accumulate at the S and G2/M phases. We concluded that Hh signalling is required for the initial entry of MuSCs into the cell cycle, and subsequently for their progression through the cell cycle.

### Hedgehog signalling controls MuSC self-renewal and long-term regenerative capability

We investigated further the role of Hh signalling in MuSC self-renewal (Figure 3E and S3A) by performing a series of three consecutive injuries at 21-day intervals in control, *Smo^cKO^* and *Ptch1^cKO^* mice (Figure 5A). Following the third round of injury, histological analyses showed the presence of a high number of mono-nucleated cells and few regenerated fibres of small size in *Smo^cKO^* mice, whereas control mice had restored a near normal muscle architecture following repeated injuries (Figure 5B,C and S5). In addition, *Smo^cKO^* muscles displayed a high degree of fibrosis (Figure 5B), a reduced number of centrally nucleated fibres (Figure 5D), and a significant increase in fibres of small size (Figure 5E) compared to control mice. This confirmed the poor regenerative capability of *Smo^cKO^* compared to control mice (Figure 5B). Together these data suggest that *Smo^cKO^* mice have poor regenerative capacity following repeated injuries, which could be due to a defect in MuSC self-renewal. Indeed, there were fewer cells able to re-integrate their sub-laminal niche in *Smo^cKO^* compared to control muscles (Figure 5F). Thus, while control mice maintain a satellite cell pool that allows them to sustain multiple injuries, *Smo^cKO^* fail to preserve a self-renewing population, leading to a poor regenerative capability upon repeated injuries.

**Figure 5.**
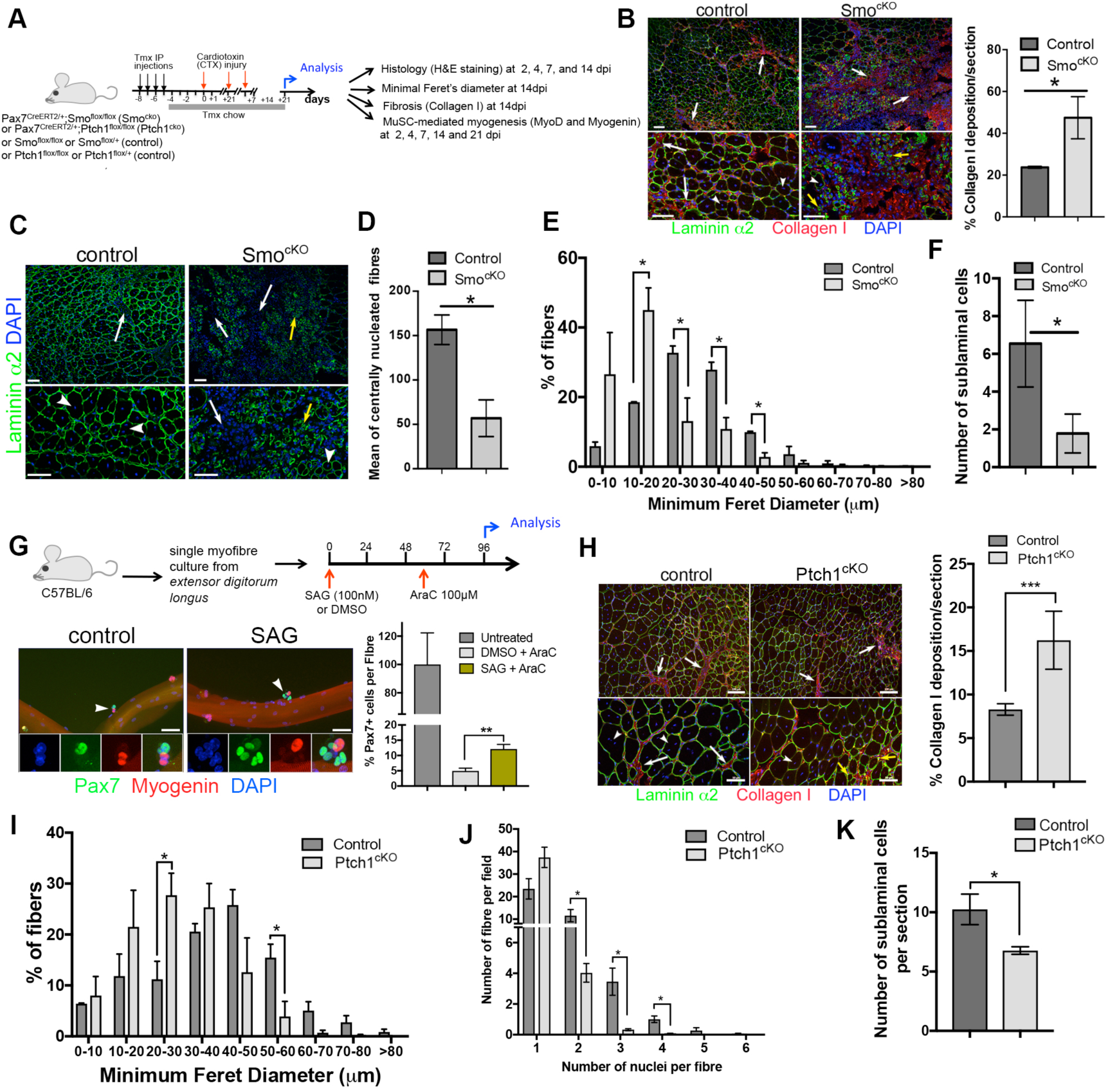
Hh signalling is necessary and sufficient to maintain a pool of self-renewing MuSCs. (A) Schematic representation of the experimental design used for (B-E). (B) Left panels: Representative images of control and *Smo^cKO^ Tibialis anterior* muscle cross-sections analysed by immunofluorescence with antibodies against Laminin α2 (green), Collagen I (red) and DAPI (blue) following three rounds of repeated injury. White arrows indicate abundant fibrosis in *Smo^cKO^* compared to control muscles. Yellow arrows indicate the presence of small regenerating fibres. White arrowheads indicate central nuclei-containing fibres. Scale bar: 50µm. Right panel: Quantification of the percentage of fibrotic area per muscle cross-section. Values are mean ± sem. unpaired t test analysis *= p < 0.05. (C) Representative images of control and *Smo^cKO^ Tibialis anterior* muscle cross-sections analysed by immunofluorescence with antibodies against Laminin α2 (green) and DAPI (blue) following three rounds of repeated injury. White arrows indicate mononucleated cells more abundant in *Smo^cKO^* than control muscles. Yellow arrows indicate the presence of small regenerating fibres. White arrowheads indicate central nuclei-containing fibres. Scale bar: 50µm (D) Quantification of the number of fibres containing centrally located nuclei in control and *Smo^cKO^* muscles. Values are mean ± sem. unpaired t test analysis *= p < 0.05. (E) Minimal Feret’s diameter analysis of control and *Smo^cKO^* muscles following three rounds of repeated injury. Values are mean ± sem. unpaired t test analysis *= p < 0.05. (F) Quantification of the number of MuSCs re-integrated in their sub-laminal niche in control and *Smo^cKO^* mice. Values are mean ± sem. unpaired t test analysis *= p < 0.05. (G) Top panel: schematic representation of experimental design. Bottom panels: representative images of immunofluorescence using antibodies against Pax7 (green) and Myogenin (red) to analyse fibers after a 96 hour-culture in the presence of DMSO (control) or SAG (100nM). Cytosine β-D-arabinofuranoside (AraC, 100µM) was added to cultures at 60 hours. White arrowhead indicates cells shown at higher magnification in inserts. Scale bar: 50µm. Right panel: Quantification of the percentage of cells positive for Pax7 after fibers were cultured for 96 hours in the absence of AraC or SAG (dark grey), or in the presence of AraC with (green) or without (light grey) SAG. Values are mean ± sem. unpaired t test analysis *= p < 0.01. (H) Left panels: Representative images of control and *Ptch1^cKO^ Tibialis anterior* muscle cross-sections analysed by immunofluorescence with antibodies against Laminin α2 (green), Collagen I (red) and DAPI (blue) following three rounds of repeated injury. White arrows indicate areas of fibrosis. Yellow arrows indicate the presence of small regenerating fibres. White arrowheads indicate central nuclei-containing fibres. Scale bar: 100µm (top images) and 50µm (bottom images). Right panel: Quantification of the percentage of fibrotic area per muscle cross-section. Values are mean ± sem. unpaired t test analysis ***= p < 0.001. (I) Minimal Feret’s diameter analysis of control and *Ptch1^cKO^* muscles following three rounds of repeated injury. Values are mean ± sem. unpaired t test analysis *= p < 0.05. (J) Quantification of the distribution of fibers per cross-section according to the number of centrally located nuclei in control and *Ptch1^cKO^* mice. Values are mean ± sem. unpaired t test analysis *= p < 0.05. (K) Quantification of the number of MuSCs re-integrated in their sub-laminal niche in control and *Ptch1^cKO^* mice. Values are mean ± sem. unpaired t test analysis *= p < 0.05.

As SAG treatments showed an increased number of Pax7^-^/MyoD^+^ cells in 72-hour muscle fibre cultures (Figure S3A) suggesting that Hh signalling can promote MuSC self-renewal, we performed single myofibre cultures in the presence of SAG or DMSO, and added the DNA replication toxin 1β-arabinofuranosylcytosine (AraC) at 60 hours when cells begin exiting the cell cycle to selectively eliminate all cycling cells. At 96 hours, myofibres were analysed for Pax7 expression to identify self-renewing cells (Figure 5G). In SAG-treated myofibre cultures, there was a significant increase in the number of self-renewing MuSCs compared to control cultures (Figure 5G), demonstrating that Hh signalling stimulates self-renewal. We then tested whether increasing levels of Hh signalling would have a beneficial effect on the long-term regenerative capability of MuSCs and performed repeated injuries in *Ptch1^cKO^* mice. However, *Ptch1^cKO^* mice did not present a noticeable improvement in their regenerative capability, and after three rounds of muscle injuries, we observed an increase in fibrosis, albeit lower than in *Smo^cKO^* mice (compare Figure 5H and 5B). Notably, although regeneration occurred, *Ptch1^cKO^* fibres were generally smaller (Figure 5I) and contained fewer nuclei per fibre (Figure 5J) compared to control fibres. In vivo, there was also a slight reduction in the number of MuSCs in a sub-laminal position, indicative of a weaker self-renewing cell population. Therefore, although increasing Hh signalling improves MuSC self-renewal in vitro and after a single muscle injury, it has no beneficial effects on the long-term regenerative capability of muscles in vivo, most likely due to additional effects on muscle progenitor cell production.

### Transcriptional profiling analyses reveal that Hh signalling controls cell cycle entry and progression

To gain further insight into the molecular and cellular processes controlled by Hh signalling in MuSCs, we performed a genome-wide transcriptome analysis of *Smo^cKO^* and *Ptch1^cKO^* MuSCs (Figure 6A). Total RNA extracted from control, Smo^cKO^ and Ptch1^cKO^ myofibres after a 62-hour culture period was subjected to next-generation RNA sequencing (RNA-seq). The analysis revealed that 306 genes were differentially expressed in *Smo^cKO^* and 211 genes were differentially expressed in *Ptch1^cKO^* compared to control samples (FDR<0.05), of which only 16 genes were common between *Smo^cKO^* and *Ptch1^cKO^* transcriptomes (Figure S6A). Comparison of control, *Smo^cKO^* and *Ptch1^cKO^* transcriptomes indicated that a number of genes down-regulated in *Smo^cKO^* were up-regulated in *Ptch1^cKO^*, and vice versa as one might expect (Figure 6B). However, the comparison of *Smo^cKO^* and *Ptch1^cKO^* transcriptomes revealed also that down-regulated *Smo^cKO^* genes were not necessarily up-regulated in the *Ptch1^cKO^* signature, as expected from gain-of-function (Figure S6B). Of interest, a number of genes associated with cell cycle regulation and muscle differentiation were down- and up-regulated in both *Smo^cKO^* and *Ptch1^cKO^* transcriptomes (Figure 6C,D). Gene ontology analysis for biological processes of the *Smo^cKO^* transcriptome indicated a transcriptional signature consistent with our observations with enriched terms for regulation of cell proliferation (GO:0042127), DNA replication (GO:0006260) and cell cycle G1/S phase transition (GO:0044843) amongst down-regulated genes and conversely enriched terms for muscle differentiation (muscle system process [GO:0003012], muscle contraction [GO:0006936] and regulation of muscle contraction [GO:0006937) amongst up-regulated genes (Figure 6E). In contrast, gene ontology analysis of the *Ptch1^cKO^* transcriptome showed an enrichment for terms associated with regulation of cell cycle (GO:0051726), regulation of cell differentiation (GO:0045595) and regulation of programmed cell death (GO:0043068) amongst up-regulated genes, whereas down-regulated genes were associated with the regulation of gene expression (GO:0010468) and gene transcription (GO:0006355), as well as the regulation of cell cycle G2/M transition (GO:1902749) and stem cell division (0017145) (Figure 6F). We validated some of the cell cycle genes identified using a multiplex bar coded fluorescent set of custom-designed probes and the Nanostring nCounter technology. This confirmed that the down-regulation of *dhx9*, a DExD/H-box nucleic acid helicase with roles in the control of transcription and of DNA replication and repair (Lee and Pelletier, 2016), of *pea-15,* a multi-functional protein known to control cell cycle progression in a ERK-dependent manner (Fiory et al., 2009), of *dck*, a kinase induced by the Ataxia-Telangiectasia Mutated (ATM) kinase following radiation-induced DNA damage and implicated in the G2/M checkpoint (Yang et al., 2012), and of *ncoa3,* a steroid receptor co-activator implicated in *cdk2* and cell cycle control (Louie et al., 2006) in the absence of Smo. Interestingly, although not statistically significant, there was down-regulation of *cdk2*, a Cyclin kinase gene associated with the control of quiescence-proliferation decision, in the absence of Smo (Spencer et al., 2013). This correlated with the significant up-regulation of *Hdac4,* a class IIa histone deacetylase whose enzymatic function is associated with muscle atrophy and satellite cell differentiation in *Smo^cKO^* muscle cells (Choi et al., 2012; Marroncelli et al., 2018). Our nCounter analysis validated also target genes significantly up-regulated in the absence of Ptch1, including *usp29,* a de-ubiquitination enzyme that controls Claspin stability and S phase progression through the Atr-Chk1 checkpoint (Martin et al., 2015). Interestingly, our data show the down-regulation of the DNA pre-replication complex gene *mcm4* and to a lesser extent that of *Dbf4,* a gene coding for the regulatory subunit of the DDK kinase controlling Mcm4 activity, both essential regulators of S phase progression in *Ptch1^cKO^* muscle cells (Sheu and Stillman, 2010). Genes implicated in the DNA damage response such as the class-IV phosphoinositide 3-kinase (PI3K)-related kinase gene *atr* and the nuclear protein coding gene *brca1* were also down-regulated in *Ptch1*-deficient MuSCs. Interestingly, both Brca1 and Atr can regulate Chk1 activity and control progression through S phase (Brown and Baltimore, 2003; Mullan et al., 2006). Thus, the analysis of *Smo^cKO^* and *Ptch1^cKO^* transcriptomes are consistent with our findings that in the absence of Hh signalling MuSCs slow-down their cell cycle and preferentially generate differentiating cells, whereas up-regulation of Hh signalling results in cells progressing more readily through the cell cycle and favouring the production of stem cells, at the expense of myogenic differentiation.

**Figure 6.**
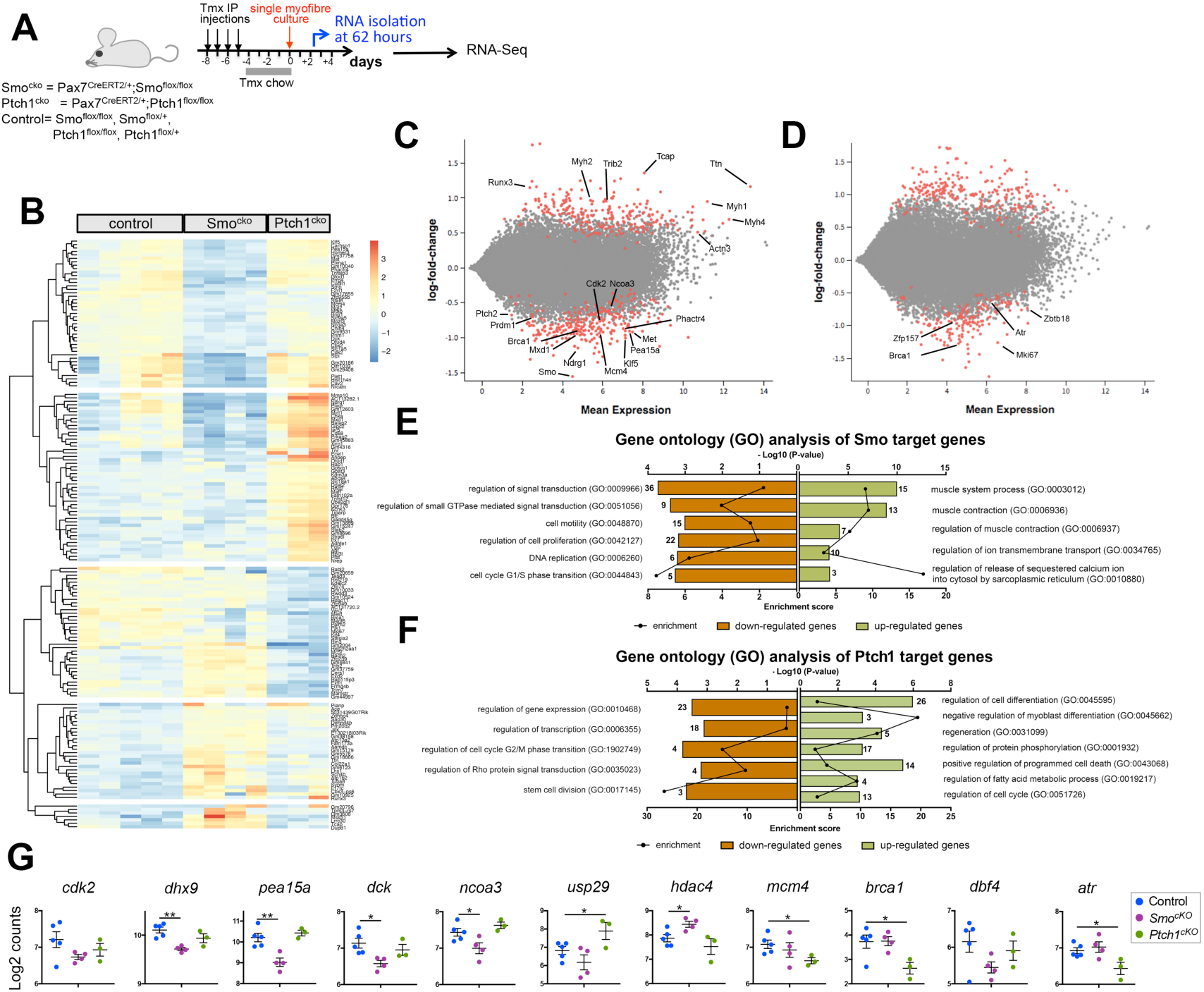
Transcriptional profiling analysis of *Smo^cKO^* and *Ptch1^cK O^* MuSCs identifies cell cycle and cell proliferation genes. (A) Schematic representation of the experimental protocol used to generate RNA for RNA-Seq analysis (B) Heatmap of some differentially expressed genes between the transcriptomes of control, Smo^cKO^ and Ptch1^cKO^ MuSCs (FDR < 0.05 and log2 fold change ±1). Columns correspond to different mice. Colors show column *Z* scores. (C) Mean average plot of differentially expressed genes in control and Smo^cKO^ MuSCs. Significantly differentially expressed genes as classified by DESeq2 are indicated in red. (D) Mean average plot of differentially expressed genes in control and Ptch1^cKO^ MuSCs. Significantly differentially expressed genes as classified by DESeq2 are indicated in red. (E) Gene ontology analysis (biological processes) of genes differentially expressed in Smo^cKO^ compared to control MuSCs. Bars represent the – log10(P value) for the term enrichment; the number of genes falling in the term enrichment category is shown above the bars; the enrichment score is indicated as a line. Enrichment terms for up-regulated genes are shown in green and enrichment terms for down-regulated genes are shown in brown. (F) Gene ontology analysis (biological processes) of genes differentially expressed in *Ptch1^cKO^* compared to control MuSCs. Bars represent the – log10(P value) for the term enrichment; the number of genes falling in the term enrichment category is shown above the bars; the enrichment score is indicated as a line. Enrichment terms for up-regulated genes are shown in green and enrichment terms for down-regulated genes are shown in brown. (G) Nanostring nCounter validation of some differentially expressed genes. Graphs represent log2(count) in control, *Smo^cKO^*, and *Ptch1^cKO^* samples. One-way ANOVA statistical analysis. *= p < 0.05; **= p < 0.01. See also Figure S6.

### The kinetics of primary cilia during myogenesis mediate a switch in Hh signalling response

Given that primary cilia are found both in quiescent MuSCs, di-assembled upon activation and selectively re-assembled in self-renewing MuSCs (Jaafar Marican et al., 2016) and that positive Hh response was observed only upon MuSC activation, we wondered whether distinct states of pathway activation could characterize Hh function in quiescent and self-renewing MuSCs. We first investigated the distribution of Smo within primary cilia in MuSCs from freshly isolated (0h) myofibres of *Tg:Pax7-nGFP* mice. In freshly isolated *Tg:Pax7-nGFP* myofibres, which contain mainly quiescent MuSCs, Smo was excluded from the primary cilium and observed in close proximity to the basal body of primary cilia (Figure 7A). We confirmed Smo association with the basal body through co-localisation with Pericentrin (Pcnt), a centrosomal protein (Figure 7B) (Jurczyk et al., 2004). Smo exclusion from the primary cilium coincided with the presence of Gpr161 within the ciliary axoneme (Figure 7A). Notably, we found that Gli3 was also localised to the basal body in quiescent MuSCs (Figure 7B). Given that Smo exclusion from the primary cilium and Gpr161 ciliary localization are associated with a repressive state of Hh signalling, which results in the processing of Gli3 by the cilia-regulated proteasome at the basal body into Gli3 repressor, our observations indicated that the primary cilium in quiescent MuSCs has important role in maintaining quiescence through the generation of Gli repressor forms. Consistent with this, the expression of *Ptch1*, a Hh target gene that is up-regulated in response to Hh signalling (Goodrich et al., 1996), increased within 12h upon myofibre culture (Figure 7C), a time that correlates with the disassembly of the primary cilium upon entry into the cell cycle (Jaafar Marican et al., 2016). To further assess this possibility, homologous recombination was induced in *Ptch1^cKO^* mice, and fibres were harvested, and either fixed immediately or cultured for 24 hours before analysis. At 0hr, *Ptch1^cKO^* MuSCs appeared to exit quiescence prematurely; a trend confirmed by 24hr when the percentage of activated MuSCs was significantly greater in *Ptch1^cKO^* fibres compared to control fibres (Figure 7D). Therefore, the primary cilium present in quiescent MuSCs mediates the repressive state of Hh signalling, and forcing Hh signalling in quiescent MuSCs causes their premature exit from G0.

**Figure 7.**
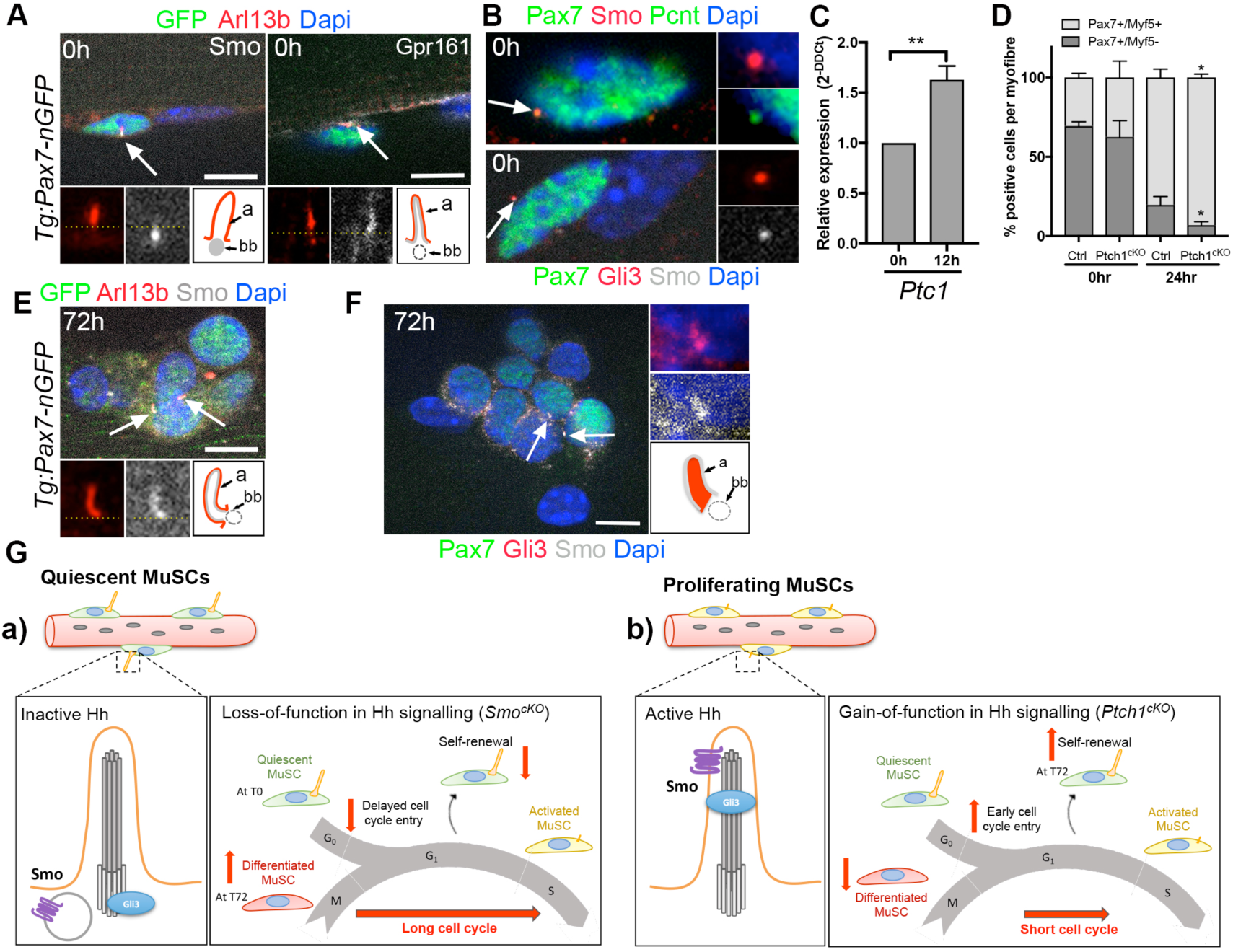
A model for Hh signalling-mediated control of MuSC quiescence, and cell cycle-dependent cell fate decision. (A) Distribution of GFP (green), Arl13b (red), Gpr161 (white), and Smo (white) in cultured Tg:Pax7-nGFP myofibres at 0h. Magnified views of red and white channels are shown with a schematic drawing of Smo and Gpr161 distribution in primary cilia. Primary cilia are indicated by white arrows. Dashed yellow line indicates the transition zone. a: axoneme; bb: basal body. Scale bar: 10µm (B) Localization of Smo and Gli3 at the cilium basal body in freshly isolated C57BL/6 muscle fibres (0hr). Top panel shows the co-localization of Smo (red) and Pericentrin (green) in Pax7^+^ MuSCs. Bottom panel shows the co-localization of Gli3 (red) and Smo (white) in Pax7^+^ MuSCs. White arrows indicate the cilium basal body. Higher magnification images are shown in inserts. (C) Relative levels of *Ptch1* mRNA expression determined by qPCR in freshly isolated (0h) and following a 12-hour culture (12h) C57BL/6 myofibres. n=4. unpaired t test analysis **= p < 0.01. (D) Graph showing the percentage of Pax7 and Myf5 positive cells per fibre in control and Ptch1^cKO^ fibres freshly isolated or cultured for 24 hours. Values are mean ± sem, n= 4 (control) and 5 (Ptch1^cKO^) with 10-24 myofibres analysed per genotype per time point. unpaired t test analysis *= p < 0.05. (E) Distribution of GFP (green), Arl13b (red) and Smo (grey) in cultured Tg:Pax7-nGFP myofibres at 72h. Magnified views of red and white channels are shown with a schematic drawing of Smo distribution in primary cilia. Primary cilia are indicated by white arrows. a: axoneme; bb: basal body. Scale bar: 10µm (F) Immunofluorescence of Pax7 (green), Smo (grey) and Gli3 (red) in cultured C57BL/6 myofibres at 72h. White arrows indicate primary cilia shown in magnified views under the main image with a schematic drawing of Smo and Gli3 distribution in primary cilia. a: axoneme; bb: basal body. Scale bar: 10µm (G) Model for Hh-mediated control of MuSC quiescence, cell cycle progression and cell fate decision. a) Maintenance of MuSC quiescence requires repressive Hh signalling through ciliary-regulated processing of Gli3 into a transcription repressor; Loss-of-function mutation in Hh signalling delays exit from quiescence whereas gain-of-function in Hh signalling promotes exit from quiescence. b) Cilia-dependent conversion of Gli3 (and possibly Gli2) into active transcription factors controls cell cycle progression in progenitor cells, and is required for MuSC self-renewal. Loss-of-function mutation in Hh signalling increases the length of cell cycle, leading to reduced self-renewal and promoting differentiation whereas gain-of-function in Hh signalling reduces the length of the cell cycle, and promotes self-renewal.

When *Tg:Pax7-nGFP* myofibres were cultured for 72h, the primary cilium re-assembled in self-renewing MuSCs. We observed that Smo relocated to the primary cilium and became uniformly distributed along the ciliary membrane of self-renewing GFP^+^ MuSCs (Figure 7E). Smo shuttling from the basal body to the primary cilium tip is associated with the transition from negative to positive transcriptional response to Hh signalling. Consistent with this, Gli3 co-localized with Smo and relocated also to the primary cilium in self-renewing Pax7^+^ MuSCs from 72h-cultured C57BL/6 myofibres (Figure 7E). Altogether, these observations suggest a switch in Hh signalling in MuSCs: in quiescent MuSCs, Hh signalling response is suppressed and this repressive state is essential to maintain MuSC quiescence. In contrast, in self-renewing MuSCs, Hh signalling is in an active state, and is required for MuSC self-renewal.

## Discussion

Skeletal muscle regenerative capacity is partly dictated by their ability to balance the use of stem cells for normal or exercise-induced growth and for repair with the need to preserve a pool of quiescent stem cells acting as a reservoir for future needs. The MuSC niche, composed of extra-cellular matrix and non-myogenic cells (Rayagiri et al., 2018; Wosczyna and Rando, 2018), provides the requisite environment to maintain this balance, and a number of extrinsic cues, including the signalling molecules Wnt, Notch and FGF and extra-cellular matrix components have previously been reported to contribute to the niche function. Here, we demonstrated that Hh signalling controls directly MuSCs, and levels of Hh signalling play a critical role in MuSC biology and muscle regeneration. We reported that the Hh signalling pathway is in a repressive state in quiescent MuSCs, and forced activation of the pathway causes MuSCs to exit quiescence prematurely. MuSCs become rapidly responsive to Hh signalling upon activation. Levels of Hh signalling are critical for MuSC cell cycle progression, as loss-of-function of Hh signalling lengthens MuSC cell cycle whereas gain-of-function of Hh signalling shortens MuSC cell cycle. Notably, Hh-regulated MuSC cell cycle progression impacts on cell fate decision upon cell cycle exit as up-regulation of Hh signalling results in increased self-renewal while down-regulation of Hh signalling causes a decline in self-renewal and concomitant increase in myogenic differentiation. We propose a model whereby the primary cilium present on quiescent MuSCs acts as a gatekeeper for Hh signalling and cell cycle entry, keeping Hh signalling in a repressive state essential for maintenance of MuSC quiescence (Figure 7G). Activation of MuSCs, which is associated with the disassembly of the primary cilium, triggers a switch in Hh signalling leading to de-repression of Hh response and entry in the cell cycle. Thereafter, levels of Hh signalling control how MuSCs progress through the cell cycle, and their decision to self-renew or differentiate upon cell cycle exit. Notably, the selective re-assembly of a primary cilium in self-renewing MuSCs allows activation of Hh signalling above de-repressed levels and drives MuSC return to quiescence (Figure 7G). Therefore, the combined activities of the primary cilium and Hh signalling provide an exquisite mechanism to control MuSC cell cycle progression and couple it to cell fate decision. These findings may have numerous implications in furthering our understanding of neuro-muscular diseases, including muscular dystrophies, muscle tumours and aging, and in designing strategies to alleviate their burden on skeletal muscles.

### Hedgehog signalling acts cell-autonomously on MuSCs

Our findings demonstrate that Hh signalling acts in a cell-autonomous manner on MuSCs, resolving a long-standing question in the field. Indeed, although up-regulation of *Ptch1* and *Gli1* has been observed in injured or diseased muscles (Piccioni et al., 2014b) and primary cilia reported at the surface of MuSCs (Fu et al., 2014; Jaafar Marican et al., 2016; Kopinke et al., 2017), previous genetic studies to inactivate or stimulate Hh signalling or to ablate primary cilia have concluded that Hh signalling acts in a non cell-autonomous manner on myogenic cells by modulating the activity of fibro-adipogenic progenitor cells, interstitial fibroblasts and immune cells (Kopinke et al., 2017). Here, we report that in addition to its role during the embryonic development of skeletal muscles (Anderson et al., 2009; Anderson et al., 2012; Borycki et al., 1999; Gustafsson et al., 2002), Hh signalling acts directly on adult MuSCs to control their activity and skeletal muscle regeneration. Such findings are likely to shed new light on the possible implication of MuSCs in the genesis of Hh-dependent rhabdomyosarcomas (Hatley et al., 2012; Hettmer et al., 2016; Rajurkar et al., 2014; Zibat et al., 2009).

### The primary cilium acts as a gatekeeper to maintain MuSC quiescence

We report that quiescent MuSCs are refractory to Hh signalling in normal adult muscle, a finding consistent with previous reports (Rajurkar et al., 2014). We showed that the primary cilium present on quiescent MuSCs is associated with a repressive state of Hh signalling, as indicated by the exclusion of Smo from the ciliary membrane and the localization of Gli3 at the basal body where it is known to be converted into a repressor form (Corbit et al., 2005; Wen et al., 2010). Pharmacological and genetic inhibition of Hh signalling delays MuSC exit from quiescence. Conversely, drug-mediated or genetic Hh signalling up-regulation causes premature MuSC exit from quiescence, indicating that cilia-mediated suppression of Hh signalling in quiescent MuSCs is an essential mechanism to maintain quiescence (Figure 7G). Interestingly, our gene profiling analysis of *Smo^cKO^* fibres identified *c-Met* as a downstream target of Hh signalling in MuSCs. Signalling from HGF to c-Met has been reported to trigger mTor signalling and promote MuSC transition from G0 to G^alert^ (Rodgers et al., 2014). Although we did not record changes in levels of *mTor* expression in our RNA-Seq analysis, it is possible that inactive Hh signalling prevents MuSCs from switching to G^alert^ by keeping *c-Met* down-regulated. Supporting our findings, recent transcriptome studies of in-situ fixed quiescent MuSCs identified genes belonging to Smo signalling pathway and intraciliary transport amongst the quiescence-enriched signature (Machado et al., 2017; van Velthoven et al., 2017). In a companion publication, Brun et al. confirmed this hypothesis by demonstrating de-repression of mTorC1 signalling in Gli3-deficient MuSCs (this issue of *Cell*).

### Levels of Hedgehog signalling are critical for MuSc cell cycle progression

Loss-of-function in Hh signalling causes an elongation of the cell cycle whereas gain-of-function in Hh signalling results in a shortening of the cell cycle in muscle progenitor cells (Figure 7G). The idea that Hh signalling acts as an internal cell cycle timer regulating the duration of the cell cycle to control the growth of developing tissues and organs is well-established in developmental biology (Chinnaiya et al., 2014; Durand et al., 1998). Our data, combined with that of others, suggests that similar mechanisms are at play in adult tissues where they control the progression of stem cells through the cell cycle, suggesting a wide-spread relationship, not initially anticipated between Hh signalling and cell cycle timing in the embryo and the adult. For instance, increased levels of Hh signalling in ventral sub-ventricular zone neural stem cells result in cell cycle shortening (Daynac et al., 2016). However, the molecular mechanisms activated downstream of Hh signalling in the control of cell cycle progression may be cell type-specific. In neural stem cells, Hh-mediated cell cycle effects involved the cyclin D-associated kinase, cdk6 (Beukelaers et al., 2011), whereas Hh regulates cell cycle timing through *cyclin D2 and p27^Kip1^* in the limb (Pickering et al., 2019). In MuSCs, Hh signalling controlled the expression of genes implicated in DNA replication and DNA damage repair, both processes tightly linked to cell cycle checkpoints and cell cycle progression (Lemmens and Lindqvist, 2019).

### MuSC cell cycle progression correlates with cell fate decision

In adult stem cells, which are normally quiescent, the rate of production of progenitor cells and cell fate decision must be tightly regulated to maintain homeostasis during the lifespan of organs. This implies a degree of coordination between cell cycle progression and decision to exit the cell cycle to self-renew or differentiate. Here, we show that loss of Hh signalling in MuSCs (*Smo^cKO^*), which causes an elongation of the cell cycle, results in a decline in self-renewing cells and a concomitant increase in myogenic differentiation (Figure 7G). In contrast, up-regulation of Hh signalling in MuSCs (*Ptch1^cKO^*), which is associated with a shortening of the cell cycle, results in an increase in the number of self-renewing cells (Figure 7G). Therefore, levels of Hh signalling control progression through the cell cycle, which in turns correlates with cell fate decision. These findings are reminiscent of studies on the developing cortex and retina showing that lengthening the cell cycle increases neurogenesis whereas shortening the cell cycle has the opposite effect (Calegari and Huttner, 2003; Lange et al., 2009). Likewise, in the developing hematopoietic lineage, the genetic ablation of *Cyclins D* delayed progression through G1, and was associated with depletion of hematopoietic stem cells (Kozar et al., 2004). In adult stem cells, the demonstration of a correlation between cell cycle progression and cell fate decision has been more challenging, and few studies have implicated Hh signalling in this regulation. However, Hh-mediated shortening of the cell cycle has been shown to transiently increase the pool of progenitor cells, causing a concomitant increase in the production of stem cells in stem cells of the ventral sub-ventricular zone, the dorsal dentate gyrus, and the liver (Daynac et al., 2016). This leads us to conclude that Hh-mediated regulation of cell cycle progression may be a common mechanism in embryonic and adult stem cells to control the balance between progenitor cell production for tissue development, growth and repair and maintenance of a stem cell pool. Clearly, future studies will need to determine the multiple expected ramifications of such principle in the control of cell polarity and cell division, metabolism, and epigenetic landscape, all processes known to be involved in cell fate decision in stem cells. The implications of such findings in basic and therapeutic research in the development, growth and repair of organs are numerous.

## Acknowledgements

We are grateful to Prof. Shahragim Tajbbakhsh, Prof. Ulrike Mayer, Prof. Giulio Cossu and Dr. Shankar Srinivas for providing us with mouse lines. We would like to thank Anna Kicheva for her technical support. The authors would like to thank Dr. Mark Dunning of the Sheffield Bioinformatics Core for assistance with RNA-seq analysis. The Sheffield Bioinformatics Core is supported / funded by the NIHR Sheffield Biomedical Research Centre (BRC) / NIHR Sheffield Clinical Research Facility (CRF). This work was funded by a grant from Association Francaise contre les Myopathies (AFM) to AGB, by a CONACYT scholarship to SBCM and by a MARA scholarship to KMI. AW and OR were recipients of Wellcome Trust summer and SURE studentships, respectively. JB is supported by the Francis Crick Institute, which receives its core funding from Cancer Research UK, the UK Medical Research Council and Wellcome Trust (all under FC001051).

## Author Contributions

SCM, KMI, JB and AGB designed the experiments. SCM, KMI, AW, OR and AGB performed the experiments. SCM, KMI, AW, OR and AGB analyzed the data. SCM, KMI, and AGB wrote and edited the manuscript.

## STAR methods

### Animal models

Mice were maintained in accordance with the UK Home Office regulation and the UK Animal Act of 1986 (ASPA) under a UK Home Office licence. All mouse lines were maintained on a C57BL/6 background and included Tg(GBS-GFP) (Balaskas et al., 2012), Tg(Pax7-EGFP) (Sambasivan et al., 2009), Pax7^tm2.1(cre/ERT2)Fan^ (thereafter named Pax7^CreERT2^) (Lepper et al., 2009), Smo^tm2Amc^ (Long et al., 2001) (thereafter named Smo^flox^), B6N.129-*Ptch1^tm1Hahn^*/J (thereafter named Ptch1flox) (Uhmann et al., 2007), Gt(ROSA)26Sor^tm1(EYFP)Cos^ (thereafter named R26R-EYFP) (Srinivas et al., 2001) and Tg(Gt(ROSA)26Sor-Fucci2) (thereafter named R26p-Fucci2) (Abe et al., 2013). Smo^flox/flox^;Pax7^CreERT2/+^ (thereafter named Smo^cKO^) and Ptch1^flox/flox^;Pax7^CreERT2/+^ mice (thereafter named Ptch1^cKO^) were generated by crossing Smo^flox/+^;Pax7^CreERT2/+^ mice with Smo^flox/fllox^ mice and Ptch1^flox/+^;Pax7^CreERT2/+^ mice with Ptch1^flox/fllox^ mice, respectively. Control animals were Smo^flox/flox^, Smo^flox/+^, ^Ptch1flox/flox^, and Ptch1^flox/+^ mice. Genotyping was performed by Polymerase Chain Reaction (PCR) on DNA extracted from ear clips using protocols reported previously (available upon request).

### Induction of Cre-mediated recombination

Tamoxifen (Tmx, Sigma-Aldrich) (10mg/ml diluted in corn oil; Sigma-Aldrich) was administered by intraperitoneal (IP) injection at a dose of 3mg/40g body weight daily for 4 days before injury in 6-8 week-old mice. Subsequently, mice were fed on a Tmx chow (approximately 40 mg/kg/day; Harlan Laboratories) for a maximum period of 14 days. Muscle injury was performed 3-5 days following the last Tmx injection.

### Muscle injury model

Mice were maintained anaesthetized with Isoflurane (IsoFlo, Abbott) and 50μl of Cardiotoxin (CTX) (10μM in PBS) from Naja mossambica (Latoxan) was injected in the left tibialis anterior (TA) muscle to induce muscle injury. Injured and contralateral (un-injured) muscles were harvested at the indicated time points after injury. Where repeated injuries were performed, three consecutive injuries were carried out by Cardiotoxin injection into the TA muscle at 21-day intervals. The minimal Feret’s diameter of fibres was determined on transverse sections immuno-labelled with Laminin α2 using the ImageJ software.

### Isolation and culture of single myofibres

Single myofibres were isolated from extensor digitorum longus (EDL) muscles following digestion by Collagenase Type I (2 mg/ml in Dulbecco’s Modified Eagle’s medium (DMEM) supplemented with Glutamax (Life Technologies) at 37°C for 80 min. Single myofibres were cultured in suspension in 60mm petri dishes coated with bovine serum albumin (BSA; Sigma-Aldrich) containing 5ml of fibre culture medium (DMEM with Glutamax, 10% horse serum (GIBCO/Invitrogen), 0.5% chick embryo extract (Seralab), and 1% Penicillin-Streptomycin-Fungizone (PSF; Sigma-Aldrich)) at 37°C and 5% CO_2_ as described previously (Collins and Zammit, 2009). When indicated, 4μM of GANT61 (Tocris Bioscience) in dimethyl sulfoxide (DMSO, Sigma-Aldrich) or 5μM of Cyclopamine in ethanol (Calbiochem) or 100nM of Smoothened Agonist (SAG, Millipore #566660) in dimethyl sulfoxide (DMSO, Sigma-Aldrich) was added to the myofibre culture medium. Concentrations were determined by adding increasing concentrations to the medium of cultured myofibres and C2C12 myoblasts, and measuring the effect on Ptch1 and Gli1 expression by RT-PCR. EdU incorporation was carried out using the Click-iT EdU assay (Invitrogen) following the manufacturer’s instructions. Briefly, myofibre cultures were incubated with 10μM EdU for 1h at 37°C prior to their harvest and subsequently fixed in 3.7% formaldehyde for 10 minutes. Fibres were washed with 3% BSA in PBS, permeabilized with 0.5% Triton 100-X in PBS and incubated with EdU reaction mix for 30 minutes in the dark. Fibres were washed with 3% BSA and blocked with 1% goat serum in PBS for 30 min and immunofluorescence staining was performed as described below. Alternatively, myofibres were transferred to Trizol (Life Technologies) if processed for qPCR.

### Immunofluorescence

Individual muscle fibres were fixed in 4% paraformaldehyde (PFA, Sigma-Aldrich) in PBS, permeabilized with 0.5% Triton 100-X (Sigma-Aldrich) in PBS and blocked for 30 minutes with 20% horse serum in PBS. Fibres were incubated overnight at 4°C with primary antibodies diluted in PBS, washed 3 times with 0.025% Tween 20 (Sigma-Aldrich) in PBS and incubated for 1h at room temperature (RT) with secondary antibodies. Finally, single fibres were washed 3 times and mounted on microscope slides in Vectashield with DAPI (Vector Laboratories) for imaging. TA skeletal muscles were fixed for 1h in 2% PFA and 0.25% Triton 100-X in PBS, washed several times in PBS and incubated overnight at 4°C in 20% sucrose (VWR Chemicals) in PBS. Muscles were then embedded in O.C.T compound and rapidly frozen in liquid nitrogen-cold isopentane (both from VWR Chemicals). Following cryosectioning (8 to 10μm on a Bright cryostat), transverse sections were blocked for 1h in 5% BSA, 1% fetal bovine serum (FBS, Life Technologies), 1% goat serum and 0.5% Triton 100-X in PBS and incubated overnight at 4°C with primary antibodies diluted in PHT (PBS with 1% heat-inactivated goat serum and 0.5% Triton 100-X). Sections were washed 3 times with PHT and incubated for 1h at RT with secondary antibodies diluted in PHT. Finally, sections were washed 3 times with PHT and then mounted in Vectashield with DAPI for imaging. Primary antibodies used in this study were as follows: mouse anti-Pax7 (1:20, Developmental Studies Hybridoma Bank, DSHB), rabbit anti-MyoD (1:50, sc-304, Santa Cruz Biotechnology, Inc.), rat anti-MyoD (1:1000, 39991 Active Motif), rabbit anti-Myf5 (1:1000, sc-302, Santa Cruz Biotechnology, Inc.), mouse anti-Myogenin (1:50, F5D, DSHB), chick anti-GFP (1:600, ab13970, Abcam), rabbit anti-Caveolin-1 (1:400, sc-894, Santa Cruz Biotechnology, Inc.), rabbit anti-Ki67 (1:300, Leica, Biosystems), rabbit anti-PH3 (1:300, MC463, Millipore), rabbit anti-Patched1 (1:50, sc-9016, Santa Cruz Biotechnology, Inc.), rabbit anti-Smoothened (1:3000, Ab38686 Abcam), goat anti-Gli3 (1:100, AF3690 R&D Systems), mouse anti-Arl13b (1:1000, 75-287 UC Davis NeuroMab facility), mouse anti-Pericentrin (1:1000, 611814 BD Biosciences), rabbit anti-Gpr161 (1:200, 13398-1-AP, Proteintech), rat anti-Laminin alpha-2 (1:300, ALX-804-190, Enzo), rabbit anti-Laminin (1:1000, Sigma-Aldrich) and rabbit anti-Collagen I (1:350, AB765, Millipore). Secondary antibodies were Alexa 488 goat anti-rabbit IgG (A11034), donkey anti-rabbit IgG (A21206), donkey anti-mouse IgG (A21202), and donkey anti-rat IgG (A21208) or Alexa 594 goat anti-rabbit IgG (A11037), donkey anti-goat IgG (A11058) and goat anti-mouse IgG (A11005) (all used at 1:500, Molecular Probes). Images were captured on a Zeiss Apotome microscope using the Axiovision imaging system. Images were assembled using Photoshop CS version 6.

### Histology

Hematoxylin and eosin staining (H&E) was performed on 4% PFA-fixed transverse sections. Frozen sections were air-dried for 1h at RT and re-hydrated for 1h with 1X PBS. Sections were then stained for 5 min in 1% eosin in methanol (Fisher Scientific) and 5 min in hematoxylin (Richard-Allan Scientific) and dehydrated through an ethanol gradient (70 to 100% in PBS). Slides were finally incubated for a short time in Xylene (Fisher Scientific) and mounted with DPX mountant (Sigma-Aldrich) for analysis.

### Quantitative PCR

Total RNA was isolated using Trizol (Invitrogen) according to the manufacturer protocol, and cDNAs were synthesized using the Superscript III First Strand Synthesis System using random hexamers (Invitrogen). qPCR was carried out on a StepOne real-time PCR instrument (Applied Biosystems) using TaqMan reagents (Applied Biosystems). The cycling conditions used were as follows: a 20 sec denaturation at 95 °C, followed by 40 cycles of 95 °C for 1 sec and 60 °C for 20 sec. Transcript levels were normalized to eukaryotic 18S rRNA (Thermo Fisher Scientific). Primers used were: Ptch1 (Mm004436026). Applied Biosystems StepOne Software V2.3 was used to analyze the data. Relative expression levels were calculated using the 2^-ΔΔCT^ method (Livak and Schmittgen, 2001).

### Statistical analyses

For comparison between two groups, unpaired two-tailed Student’s t test was performed to calculate p values and to determine statistically significant differences (*, p < 0.05; **, p < 0.01; ***, p < 0.001, ****, p < 0.0001). For comparison between three groups, one way ANOVA test was performed. All statistical analyses were performed using the GraphPad PRISM software. Errors bars on graphs are standard errors of the mean (sem).

### Next-generation RNA sequencing (RNA-seq)

Samples analysed by RNA-seq were total RNA prepared from EDL myofibres isolated from control (n=5), Smo^cKO^ (n=4) and Ptch1^cKO^ (n=3) mice and cultured for 62 hours. Total RNA was extracted using the Nucleospin RNA plus XS purification kit (Macherey-Nagel) according to the manufacturer protocol. RNA purity was determined on a Agilent Bioanalyser 2100, and the samples RIN ranged between 9.40 and 9.70. Approximately 200ng of total RNA underwent rRNA depletion using the NEB rRNA depletion kit (catalogue number E6310). Sequencing libraries were prepared using the NEB Ultra II Directional RNA Kit for Illumina (catalogue number E7760). Raw fastq files were processed using the bcbio workflow (https://bcbio-nextgen.readthedocs.io/en/latest/). Specifically, the alignment-free quantification method salmon (Patro et al., 2017) was used to obtain counts for all transcripts in mouse gencode version 18 (genome version GRCm38.6). The tximport Bioconductor package (Soneson et al., 2015) was then used to create gene-level counts for analysis. Exploration of the quality assessment and differential expression analysis was facilitated by the bcbioRNASeq Bioconductor package (Steinbaugh et al., 2017). Ranked differential expression test statistics as calculated by DESeq2 (Love et al., 2014) were used as input to fgsea (Korotkevich et al., 2019) to perform pathway enrichment analysis.

### Nanostring nCounter analysis of gene expression

A custom Element TagSet of capture- and reporter probes was designed to target regions of some differentially expressed genes identified in the RNA-Seq analysis. Approximately 200 ng of total RNA from each sample was subjected to nCounter™ SPRINT (NanoString Technologies, Seattle, WA, USA) analysis according to the manufacturer’s instructions. The raw data were processed using the nSOLVER 4.0 software (NanoString Technologies); The data was then exported to Excel and Prism 7.0 for statistical analysis and graph production.

