## Supplementary material for "A switch in cilia-mediated Hedgehog signaling controls muscle stem cell quiescence and cell cycle progression": Figure S

### Supplemental data

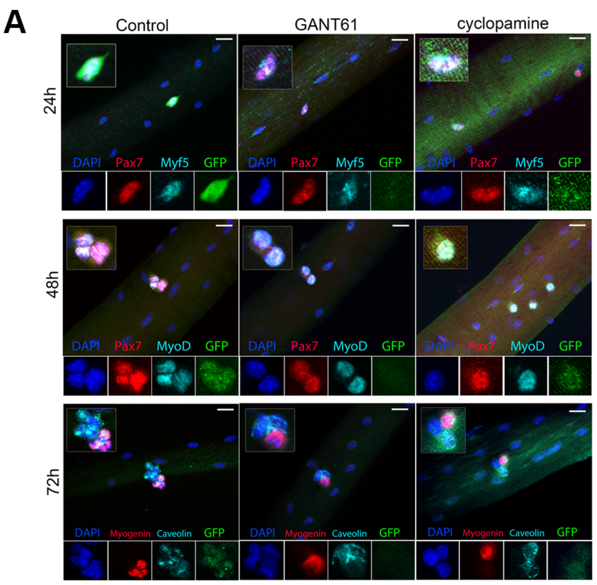


**Figure S1: MuSC Hedgehog response is down-regulated upon treatment with GANT61 or cyclopamine**

(A) Representative images of isolated Tg(GBS-GFP) myofibres cultured for 72 hours in the presence of DMSO (control), GANT61 (4µM) or Cyclopamine (5µM) and analysed by immunofluorescence using antibodies against GFP (green), Pax7 (red), Myf5 (Cyan), MyoD (cyan), Myogenin (red) and Caveolin-1 (cyan). Nuclei were counterstain with DAPI. Inserts show higher magnification merge images. Individual channel images are shown below the main panel. Scale bar: 20µm

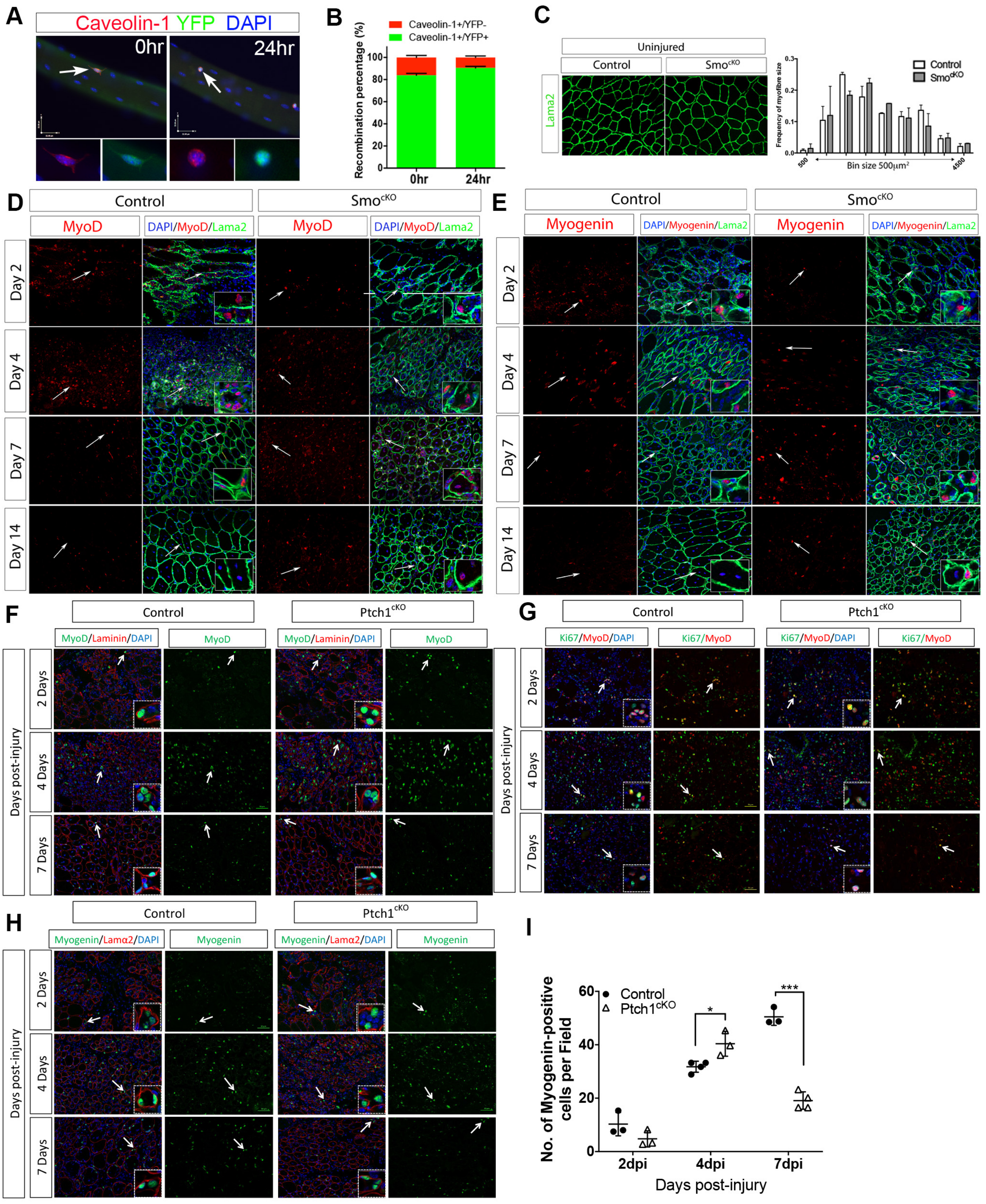


**Figure S2: Skeletal muscle regeneration following injury in *Smo*$^{cKO}$ and *Ptch1*$^{cKO}$ mice.**

(A) Representative images of freshly isolated or 24 hour-cultured myofibres from *Pax7*$^{CreERT2/+}$*; R26YFP;Smo*$^{flox/flox}$ mice analysed by immunofluorescence using antibodies against GFP (green) and Caveolin-1 (red). Nuclei were counterstained with DAPI. White arrows indicate cells shown at higher magnification below the main panel.

(B) Graph showing the percentage of recombination ($YFP^+$ cells, green bars) in freshly isolated or 24 hour-cultured myofibres from *Pax7*$^{CreERT2/+}$*; R26YFP;Smo*$^{flox/flox}$ mice. n= 3 with a total of 50 fibres analysed per time point.

(C) Left panel: representative images of un-injured control and *Smo*$^{cKO}$ TA muscles analysed by immunofluorescence with anti Laminin α2 antibodies (green). Right panel: Fibre size distribution (minimal Feret's diameter) of control and *Smo*$^{cKO}$ TA muscles. No statistical difference is observed between control and *Smo*$^{cKO}$ TA muscles. n=3.

(D) Immunofluorescence analysis of MyoD distribution (red) in control and *Smo*$^{cKO}$ TA muscles between 2dpi and 14dpi. Fibre basal lamina is labelled with Laminin α2 antibodies (green). Nuclei are counterstained with DAPI. White arrows indicate cells shown as high magnification in inserts.

(E) Immunofluorescence analysis of Myogenin distribution (red) in control and *Smo*$^{cKO}$ TA muscles between 2dpi and 14dpi. Fibre basal lamina is labelled with Laminin α2 antibodies (green). Nuclei are counterstained with DAPI. White arrows indicate cells shown as high magnification in inserts.

(F) Immunofluorescence analysis of MyoD distribution (green) in control and *Ptch1*$^{cKO}$ TA muscles between 2dpi and 7dpi. Fibre basal lamina is labelled with Laminin α2 antibodies (red). Nuclei are counterstained with DAPI. White arrows indicate cells shown as high magnification in inserts.

(G) Immunofluorescence analysis of MyoD (red) and Ki67 (green) expression in control and *Ptch1*$^{cKO}$ TA muscles between 2dpi and 7dpi. Nuclei are counterstained with DAPI. White arrows indicate cells shown as high magnification in inserts.

(H) Immunofluorescence analysis of Myogenin distribution (green) in control and *Ptch1*$^{cKO}$ TA muscles between 2dpi and 7dpi. Fibre basal lamina is labelled with Laminin α2 antibodies (red). Nuclei are counterstained with DAPI. White arrows indicate cells shown as high magnification in inserts.

(I) Quantification of Myogenin-expressing cells in control and *Ptch1*$^{cKO}$ TA muscles between 2dpi and 7dpi. n=3-4 mice per genotype. Values are mean ± sem. unpaired t test analysis *= $p < 0.05$; ***= $p < 0.001$.

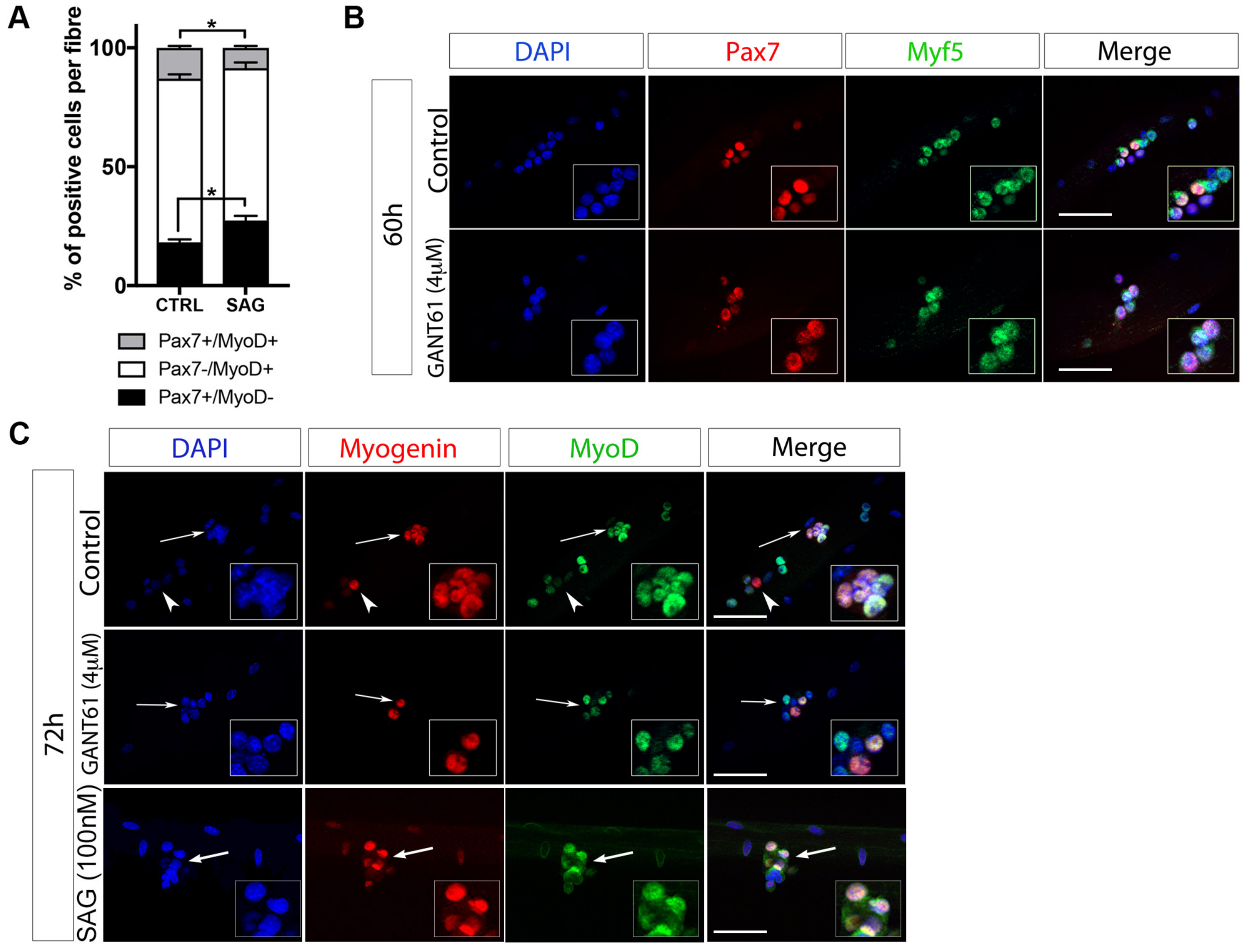


**Figure S3: Levels of Hedgehog signalling control MuSC progression through the myogenic programme.**

(A) Graph showing the percentage of Pax7 and MyoD positive cells per myofibre after a 72-hour culture period in the presence of DMSO (control) or SAG (100nM). Values are mean ± sem. unpaired t test analysis *= $p < 0.05$.

(B) Representative images of myofibres cultured for 60 hours in the presence of DMSO or GANT61 (4µM), and analysed by immunofluorescence using antibodies against Myf5 (green) and Pax7 (red). Nuclei are counterstained with DAPI (blue). Inserts show high magnification images. Scale bar: 50µm.

(C) Representative images of myofibres cultured for 72 hours in the presence of DMSO, or GANT61 (4µM), or SAG (100nM), and analysed by immunofluorescence using antibodies against MyoD (green) and Myogenin (red). Nuclei are counterstained with DAPI (blue). White arrows indicate cells shown at high magnification in inserts. Scale bar: 50µm.

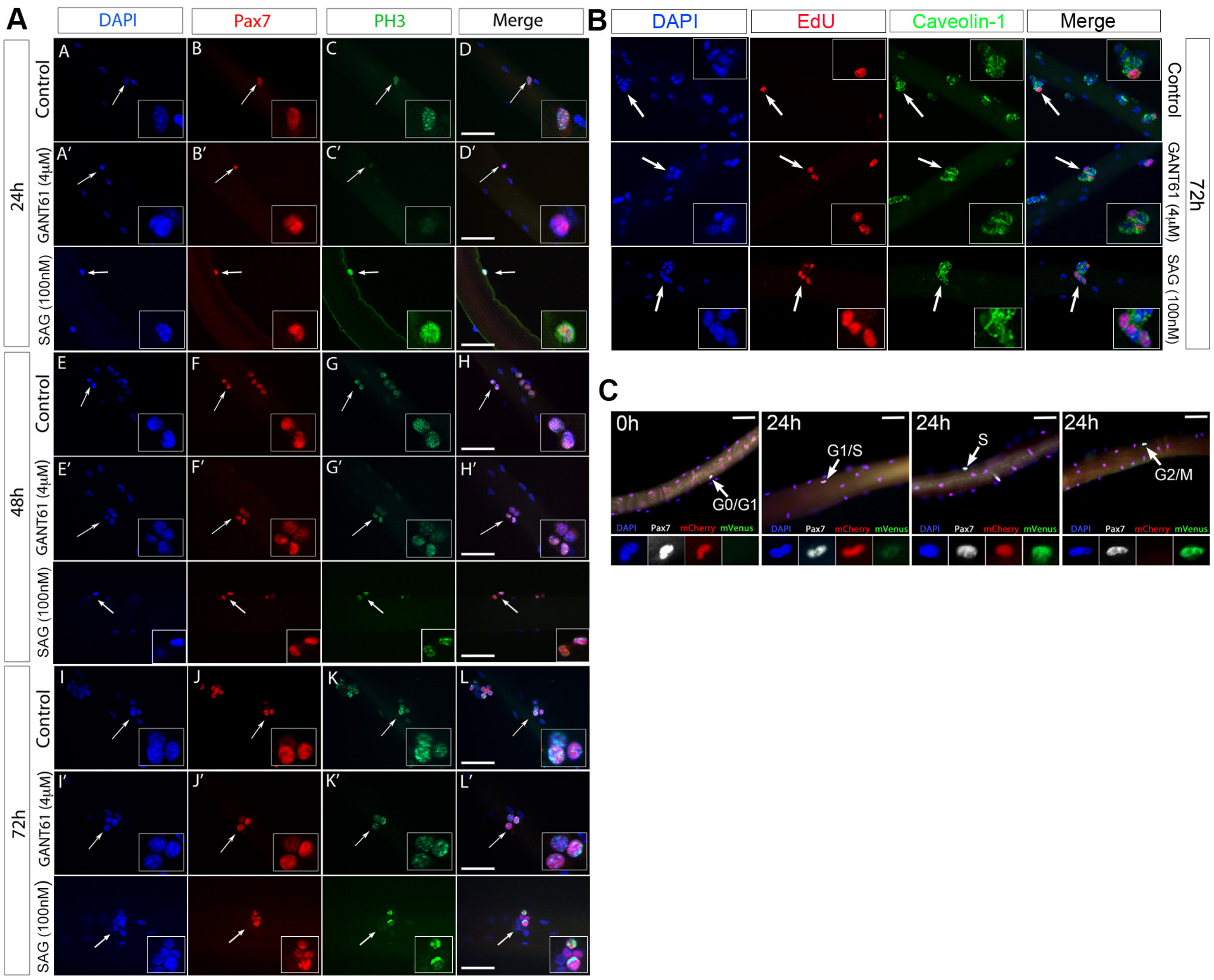


**Figure S4: Levels of Hedgehog signalling control MuSC cell cycle progression**

(A) Representative images of C57BL/6 myofibres cultured for 72 hours in the presence of DMSO or GANT61 (4μM) or SAG (100nM), and analysed by immunofluorescence using antibodies against phosphor histone H3 (PH3, green) and Pax7 (red). Nuclei are counterstained with DAPI (blue). White arrows indicate cells shown at high magnification in inserts. Scale bar: 50μm.

(B) Representative images of C57BL/6 myofibres cultured for 72 hours in the presence of DMSO or GANT61 (4μM) or SAG (100nM). EdU (red) was added to the culture medium one hour prior to the end of culture. Myofibres were analysed by immunofluorescence using antibodies against Caveolin-1 (green). Nuclei are counterstained with DAPI (blue). White arrows indicate cells shown at high magnification in inserts.

(C) Representative images of Tg(Fucci2) myofibres freshly isolated or cultured for 24 hours to illustrate the detection of G0/G1, G1/S, S, and G2/M phases. Fibres were analysed by immunofluorescence using antibodies against Pax7 (white) and GFP (green). mCherry was directly detected. White arrows indicate cells shown at high magnification in panels below the main image. Scale bar: 50μm

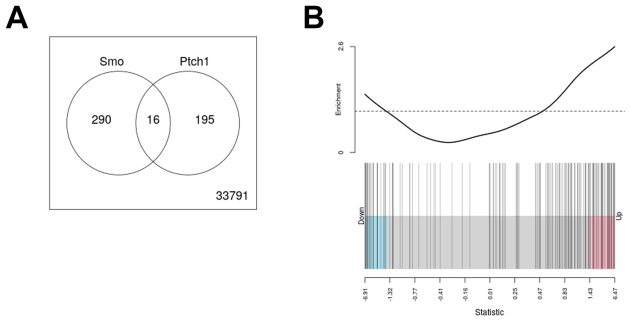


**Figure S6: Analysis of RNA-Seq data**

(A) Venn diagram of differentially expressed genes in *Smo*$^{cKO}$ and *Ptch1*$^{cKO}$ transcriptomes.

(B) Barcode plot showing the enrichment of *Smo*$^{cKO}$ down-regulated genes in the comparison of the *Smo*$^{cKO}$ vs *Ptch1*$^{cKO}$ signatures. Red bars show up signature genes, blue bars show down genes. Genes down-regulated in the *Smo*$^{cKO}$ signature tend to be up or down-regulated in the *Ptch1*$^{cKO}$ signature.
